# High-resolution profiling reveals coupled transcriptional and translational regulation of transgenes

**DOI:** 10.1101/2024.11.26.625483

**Authors:** Emma L. Peterman, Deon S. Ploessl, Kasey S. Love, Valeria Sanabria, Rachel F. Daniels, Christopher P. Johnstone, Diya R. Godavarti, Sneha R. Kabaria, Conrad G. Oakes, Athma A. Pai, Kate E. Galloway

## Abstract

Concentrations of RNAs and proteins provide important determinants of cell fate. Robust gene circuit design requires an understanding of how the combined actions of individual genetic components influence both mRNA and protein levels. Here, we simultaneously measure mRNA and protein levels in single cells using HCR Flow-FISH for a set of commonly used synthetic promoters. We find that promoters generate differences in both the mRNA abundance and the effective translation rate of these transcripts. Stronger promoters not only transcribe more RNA but also show higher effective translation rates. While the strength of the promoter is largely preserved upon genome integration with identical elements, the choice of polyadenylation signal and coding sequence can generate large differences in the profiles of the mRNAs and proteins. We used long-read direct RNA sequencing to characterize full-length mRNA isoforms and observe remarkable uniformity of mRNA isoforms from the transgenic units. Together, our high-resolution profiling of transgenic mRNAs and proteins offers insight into the impact of common synthetic genetic components on transcriptional and translational mechanisms. By developing a novel framework for quantifying expression profiles of transgenes, we have established a system for comparing native and synthetic gene regulation and for building more robust transgenic systems.

## Introduction

Intracellular levels of key proteins and RNAs govern gene regulatory programs and cell states. Similarly, levels of RNA and protein components can set the activity of gene circuits and influence the robustness and performance of the circuits. Eukaryotic gene expression requires multiple co- and post-transcriptional processing steps of mRNA transcripts including splicing, 3’ end cleavage and polyadenylation, and nuclear export [1]. The degree of post-transcriptional or post-translational processing influences the correlation between mRNA and protein levels [2]. This multi-scale process generates endogenous mRNA and protein levels that are much less correlated in eukaryotes than in prokaryotes [3]. While the number of studies profiling the expression of endogenous RNA transcripts is rapidly growing [4, 5], there is still limited understanding of the abundance and composition of mRNA expressed from synthetic transgenic systems. Developing a predictive understanding of the levels of mRNA isoforms and their rates of processing may improve the design of gene circuits in diverse cell types—including primary cells and induced pluripotent stem cells (iPSCs) [6–10].

Forward design of gene circuits requires composable and well-characterized genetic elements. Given the distribution of gene expression profiles from transgenes, models that accurately predict the performance of dynamic circuits require parameters that capture the ensemble features such as the mean and variance of mRNA and protein molecules. Accurate estimation of the average level of protein expression can inform selection of genetic elements for predictive design [11]. Tracking distributions of both transgenic mRNAs and proteins over time can augment the design of systems that amplify or attenuate noise and reveal the underlying network structures of biological systems [12–14]. However, synthetic parts are often characterized by the mean level of expression for a single mRNA or protein species. The complex nature of gene regulation in mammalian cells calls for high-resolution, systematic characterization of genetic parts across transcriptional and translational processes. Defining the combined effects of genetic elements, such as promoters, polyadenylation signals (PASs), and untranslated regions (UTRs) on gene expression profiles and transcript isoforms will offer insight into sources of variability. For instance, are mRNA isoforms uniform within a single construct? Or do transgenes generate a variety of isoforms that may have unique processing rates? Understanding both mRNA levels and compositions will support improved design of transgenic systems in mammalian cells.

As multi-modal circuits rely on levels of both RNAs and proteins, predictable circuit design requires high-resolution characterization of both molecules and their distributions across populations. Previous characterizations of genetic parts relied on easily assayable metrics of expression, such as protein fluorescence or enzymatic activity, which cannot capture species involved in post-transcriptional regulation such as microRNAs, alternative splicing isoforms, and ribozymes [2, 3, 11, 15–17] (Figure 1A). Forward design of RNA-based control systems requires quantification of RNA levels in single cells [18]. Bulk methods such as reverse transcription quantitative polymerase chain reaction (RT-qPCR) obscure variance across single cells [19]. While transcriptional imaging systems and single molecule fluorescence *in situ* hybridization (smFISH) enable high-resolution quantification of transcript profiles in single cells, these methods suffer from being very low-throughput [20–23]. Alternatively, flow cytometry-based RNA fluorescence *in situ* hybridization (Flow-FISH) offers a method to measure levels of specific RNAs in single cells. Flow-FISH enables high-throughput, sensitive RNA readouts that can be coupled with simultaneous protein quantification in a single cell. Thus, Flow-FISH allows us to measure single-cell mRNA distributions while integrating existing protein expression analysis pipelines for a more comprehensive characterization of existing and novel genetic elements. Flow-FISH paired with methods to analyze full-length isoforms would enable an understanding of how genetic elements influence variability in transcriptional and translational processes.

**Figure 1.**
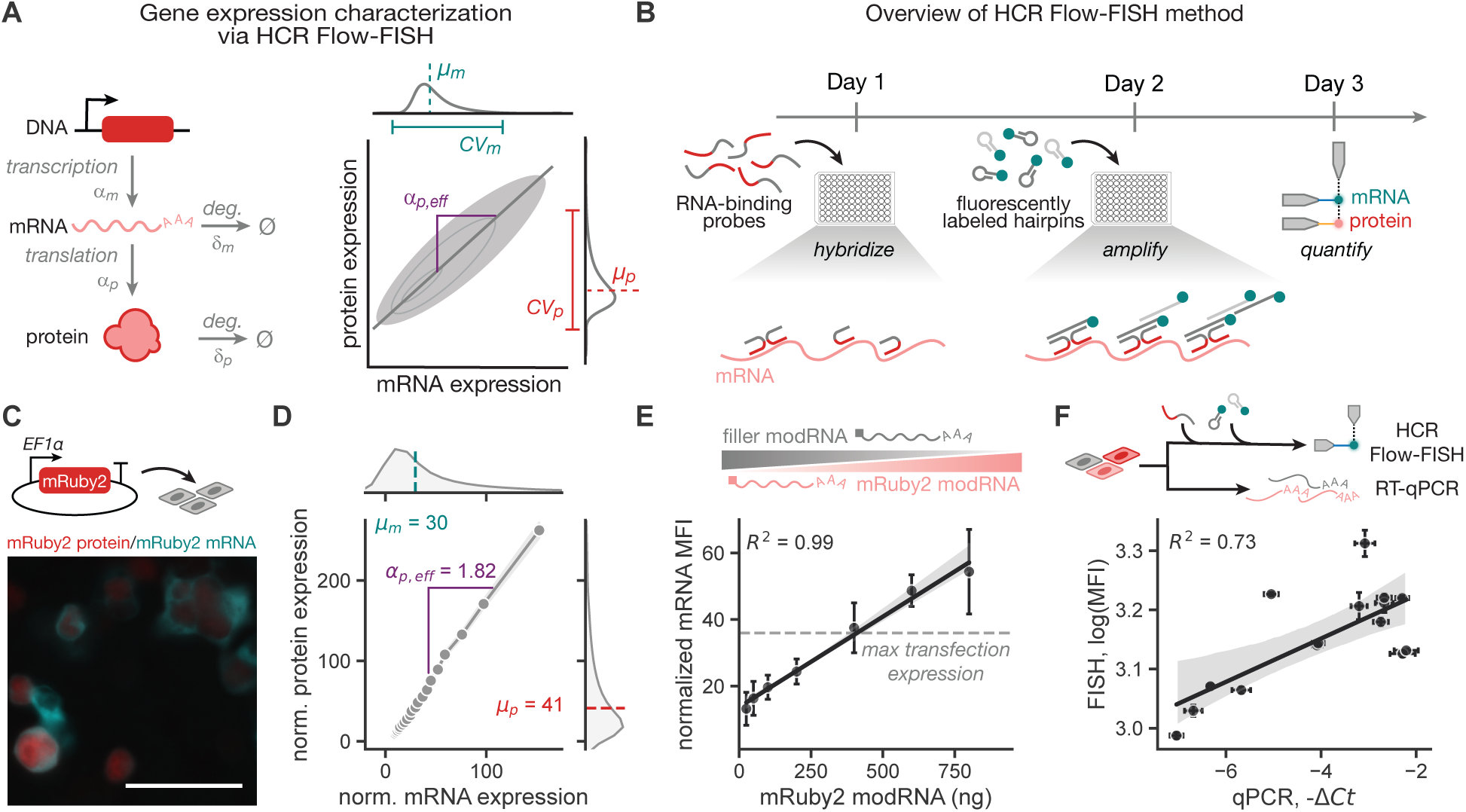
HCR Flow-FISH enables high-throughput quantification of mRNA and protein levels in single cells via flow cytometry. A. In a simple model of gene expression, mRNA and protein levels (*µ*_m_ and *µ*_p_) are governed by four main parameters: transcription rate (*α*_m_), mRNA degradation rate (*δ*_m_), translation rate (*α*_p_), and protein degradation rate (*δ*_p_). HCR Flow-FISH data allows for calculation of an effective translation rate, *α*_p,eff_ = Δ*µ*_p_*/*Δ*µ*_m_ ∝ *α*_p_*/δ*_p_, and coefficient of variation (CV). B. RNA-binding probes specific to the mRNA species of interest are first added to fixed and permeabilized cells. Fluorescently labeled hairpins complementary to the probes are then added to amplify the FISH signal. **C**, **D**. Sample HCR FISH imaging (**C**) and flow cytometry (**D**) for HEK293T cells transfected with an EF1_α_-mRuby2-bGH plasmid and labeled with Alexa Fluor™ 514 HCR amplifiers. Scale bar represents 50 µm. Data for one representative biological replicate is binned by transfection marker level into 20 equal-quantile groups. Points represent geometric mean (MFI) of protein and mRNA expression for cells in each bin, and shaded regions represent the 95% confidence interval. Effective translation rate is calculated as the slope of a line fitted to the binned data (*R*^2^ = 0.999). Normalized expression is calculated as the fold change of fluorescence intensity relative to a non-transfected sample. **E**. Measurement of HCR Flow-FISH signal for HEK293T cells transfected with varying dosages of mRuby2 modRNA. Error bars represent the standard deviation across three biological replicates, and the shaded region represents the bootstrapped 95% confidence interval of the linear regression (*p* = 7 × 10*^−^*^6^). Grey dashed line indicates the mean mRNA MFI for the highest expressing construct (CAG-mRuby2-bGH) in HEK293T transfection. Normalized fluorescence is calculated as the fold change of fluorescence intensity relative to a non-transfected sample. **F**. Measurement of mRuby2 mRNA level via HCR Flow-FISH (y-axis) and RT-qPCR (x-axis) for PiggyBac-integrated HEK293T cells with varying levels of mRuby2 expression. Error bars represent the standard deviation between technical replicates, and the shaded region represents the bootstrapped 95% confidence interval of the linear regression (*p* = 2 × 10*^−^*^3^). Each point represents an individual biological replicate. All HCR Flow-FISH data is in arbitrary units from a flow cytometer.

In this work, we use Flow-FISH to simultaneously quantify levels of transgenic mRNAs and proteins in single cells. With these data, we can quantify the impact of individual genetic elements on different gene regulatory steps. Specifically, we characterize a panel of commonly used constitutive promoter sequences in HEK293T cells and benchmark expression levels against three inducible promoter systems (Tet-On, COMET [24], and synZiFTR [25]). We find that promoter sequences impact both the abundance of mRNA transcripts and the effective translation rate of these transcripts. Moreover, the combination of promoter, coding, and 3’ UTR sequences alter the effective translation rate, suggesting a role for UTRs in transgene regulation. To examine RNA isoforms—including their UTRs—at high resolution, we use long-read sequencing to profile full-length transcripts from transgenes. We find that mature transgenic transcripts are highly uniform, rarely impacted by local sequence context, and exert minimal burden on endogenous gene expression. Together, our work to establish high-resolution profiling of expression distributions and isoforms of transgenic mRNAs offers a novel framework for systematically comparing native and synthetic gene regulation and building more robust transgenic systems.

## Results

### HCR Flow-FISH enables simultaneous, high-throughput quantification of mRNA and protein levels in single cells

To simultaneously assess mRNA and protein distributions, we used a flow cytometry-based RNA fluorescence *in situ* hybridization (Flow-FISH) method, which allows for the concurrent measurement of mRNA and protein levels in single cells [26]. Hybridization chain reaction RNA-FISH (HCR Flow-FISH) amplifies signal, improving the signal-to-noise ratio and enabling better mRNA detection at low expression levels [27, 28]. HCR Flow-FISH uses a two-stage amplification approach (Figure 1B). First, RNA-binding probes complementary to the mRNA transcript of interest are hybridized overnight in fixed and permeabilized cells. The following day, probe-specific, fluorescently labeled hairpins are added to amplify the FISH signal. HCR leads to higher fluorescence levels while minimizing background, making it well-suited for flow cytometry quantification.

We optimized and validated an HCR Flow-FISH protocol based on methods reported in Choi *et al.* [28]. We transfected HEK293T cells with a plasmid expressing the fluorescent protein mRuby2 with the EF1α promoter and bGH PAS (EF1α-mRuby2-bGH). We quantified expression of the mRuby2 mRNA using sequence-specific FISH probes and compatible Alexa Fluor™ 514 HCR amplifiers. Through fluorescence imaging (Figure 1C) and flow cytometry (Figure 1D), we measured both mRNA and protein expression in the same single cells. Binning cells based on expression of a co-transfected fluorescent marker, we observe a linear dependence between mRNA signal and protein signal. We quantified a dimensionless effective translation rate of the transcripts, *α*_p,eff_ by calculating the slope between mRNA signal and protein signal via least squares regression (Figure 1A). This parameter is proportional to the number of proteins translated from a single mRNA transcript and combines the contributions of mRNA transport, translation initiation, and protein stability.

To verify that HCR Flow-FISH detects quantitative changes in mRNA level, we transfected varying dosages of mRuby2-encoding modRNA into HEK293T cells. To minimize any differences in transfection efficiency, we added a non-fluorescent, “filler” modRNA to maintain a consistent total modRNA amount across all conditions. As expected, the mean HCR Flow-FISH signal increases linearly with modRNA dosage, indicating that this method can detect the anticipated mean differences in mRNA levels between populations of cells (Figure 1E). Importantly, when transfecting a plasmid with a strong constitutive promoter, CAG (dashed line), the measured HCR Flow-FISH signal sits within the linear detection regime and does not saturate. Compared to genomic integration, transfection typically leads to higher transgene DNA copy numbers, so this condition represents the maximum relevant amount of mRNA for detection. Therefore, because HCR Flow-FISH signal correlates linearly with mRNA level across the range of expression levels, we can quantitatively compare mRNA levels between different compositions of genetic elements. Additionally, we find that the mean signal from HCR Flow-FISH correlates positively with the signal from RT-qPCR across five cell lines with varying mRuby2 expression (Figure 1F). Thus, we conclude that HCR Flow-FISH and RT-qPCR provide similar relative estimates for mean mRNA levels, while HCR Flow-FISH offers the additional benefit of quantifying the distribution of mRNA levels. Together, HCR Flow-FISH enables single-cell quantification of mRNA levels for simultaneous characterization of mRNA and protein expression distributions.

### Promoter sequences affect RNA transcript abundance and effective translation rate

With the quantitative HCR Flow-FISH protocol validated, we characterized a commonly used set of genetic elements including promoters. In the field of synthetic biology, genetic components that drive transcription are determined heuristically from native genomes and often include the combination of a minimal promoter, a transcription start site, upstream regulatory elements, and in some cases an associated 5’ UTR sequence. In this work, we refer to this entire set of sequences as the promoter. Choice of promoter can set the level of protein expression. However, as promoters differ in recruitment of transcriptional machinery, 5’ UTR sequences, and splicing within 5’ UTRs, protein data alone cannot define how promoters influence transcription rates. We chose to evaluate a set of six constitutive promoters with varying expression levels, composition, and origins: CAG, a hybrid of the cytomegalovirus enhancer and chicken beta-actin promoter; EF1α, the human elongation factor 1-alpha promoter; CMV, a strong promoter derived from cytomegalovirus; UbC, the human polyubiquitin C promoter; hPGK, the human phosphoglycerate kinase promoter; and EFS, a derivative of the EF1α promoter lacking the intron (Table S1). To assess the relative expression levels of this promoter set, we used each promoter to drive expression of mRuby2 with a bGH PAS (Figure 2). For each condition, we included a co-transfected marker plasmid encoding a separate fluorescent protein, whose expression we used as a proxy for copy number [29]. Constitutive promoter conditions were analyzed via HCR Flow-FISH at two days post-transfection (Figure 2A). As expected from previous characterizations [15], our data ranks promoters by strength— as measured by mean protein expression—from highest to lowest: CAG, EF1α, CMV, UbC, EFS, and hPGK (Figure S1).

**Figure 2.**
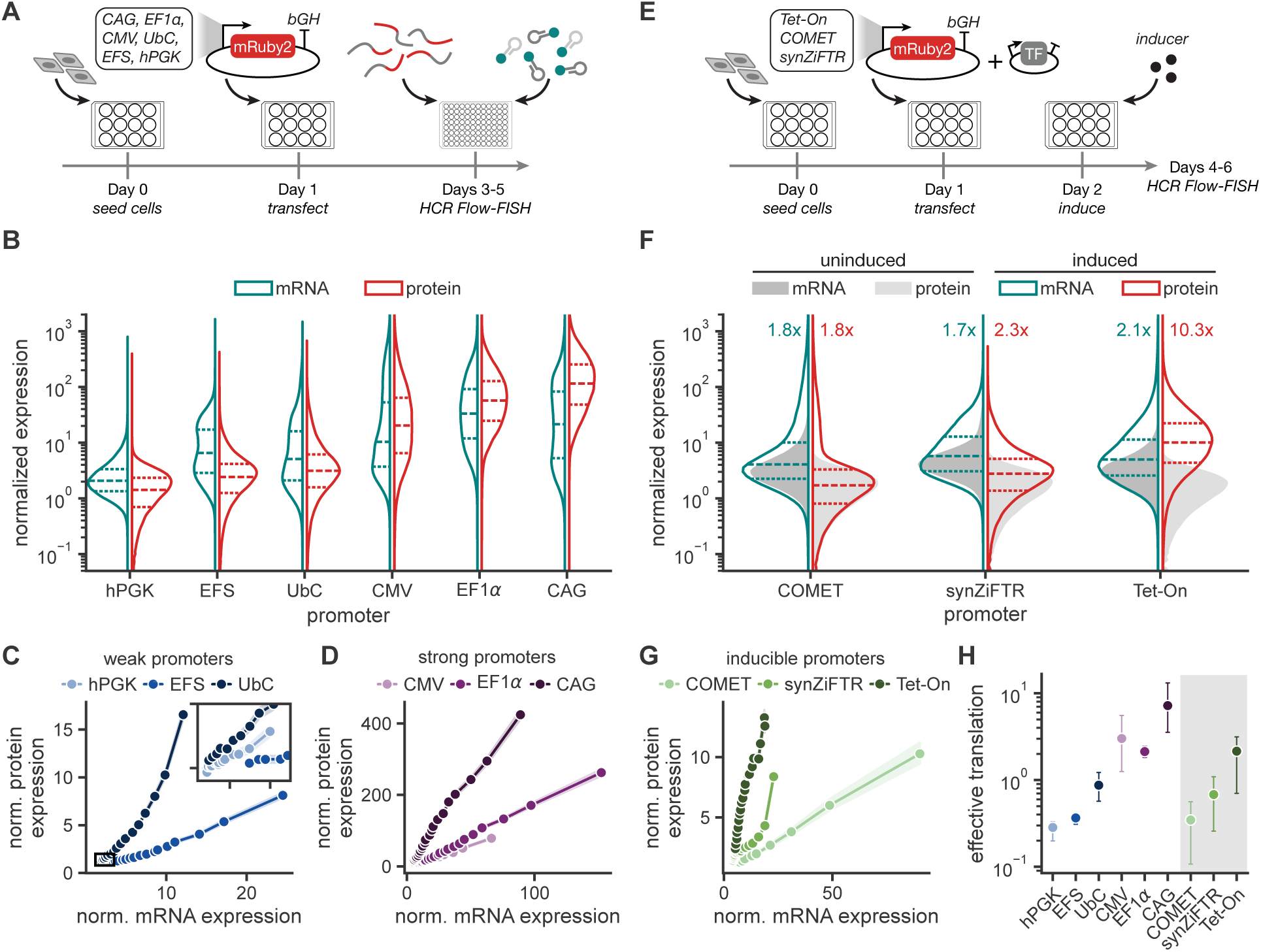
Promoter sequences affect RNA transcript abundance and effective translation rate. **A.** HEK293T cells were transfected with a plasmid encoding mRuby2 driven by one of six different constitutive promoters (CAG, EF1_α_, CMV, UbC, EFS, or hPGK). **B.** Normalized expression distributions for mRNA (Alexa Fluor™ 514) and protein (mRuby2) with six constitutive promoters as measured by flow cytometry. **C.** Normalized protein expression as a function of normalized mRNA expression with three weak constitutive promoters. Inset axes show the indicated low expression domain. Data for one representative biological replicate is binned by marker level into 20 equal-quantile groups. Points represent geometric means of mRNA and protein levels for each bins. Shaded regions represent the 95% confidence interval. **D.** Normalized protein expression as a function of normalized mRNA expression for three strong constitutive promoters for one representative biological replicate. **E.** HEK293T cells were transfected with plasmids encoding mRuby2 driven by one of three different inducible promoters (Tet-On, COMET, synZiFTR) and the corresponding transcription factor. **F.** Normalized expression distributions for mRNA (Alexa Fluor™ 514) and protein (mRuby2) with three inducible promoters following small molecule induction as measured by flow cytometry. Fold-change in expression upon induction is annotated for each condition. **G.** Normalized protein expression as a function of normalized mRNA expression with three inducible promoters for one representative biological replicate. **H.** Effective translation rate as calculated by the slope of a line fitted to the binned data. Inducible promoters are denoted by the gray box. Points represent the mean of three biological replicates ± the 95% confidence interval. Normalized expression is calculated as the fold change of fluorescence intensity relative to a non-transfected sample. All data is in arbitrary units from a flow cytometer.

In transfection, mRNA expression exhibits more variance across the population than protein expression (Figures S1 and 2B). Bimodality observed in levels of mRNA and not in protein may reflect the higher stability of the proteins compared to mRNA. Despite the increased variance, the relative ordering of promoter strength, as determined by the mean mRNA expression level, generally matches that of the mean protein expression level (Figure S1). However, CAG achieves the highest protein expression but has only the second-highest level of mRNA, indicating post-transcriptional processing and translation rates influence protein levels. Binning the cells by expression of the transfection marker, we find that higher mRNA levels are strongly correlated with higher protein levels (Figures 2C and 2D). However, the relative slopes of these curves differ, indicating differences in effective translation rate across promoters. In addition to having higher mRNA levels, the strong promoters (CAG, EF1α, and CMV) exhibit more efficient translation than the weak promoters (UbC, EFS, and hPGK) (Figure 2H). These differences are likely sequence-specific and could be attributed to factors such as RNA nuclear export, localization, and secondary structure.

To assess generality of trends, we characterized profiles of expression across different cell types and integration methods. First, we quantified profiles of expression for transfection in Chinese hamster ovary (CHO-K1) and iPS11 cells. Both cell types maintain the same relative promoter strengths, with the exception of CMV, which did not express above background in iPS11 cells (Figure S2A). Immunogenicity associated with viral sequences in CAG and CMV may suppress gene expression in iPSCs [30]. We next explored how integration into the genome affects profiles of expression. Understanding how transfection profiles translate to integration can accelerate the design-build-test-learn loop, which remains slower due to the time scales of generating cell lines. We quantified expression from each promoter in HEK293T cells after random integration via PiggyBac transposase and lentivirus as well as site-specific integration at a landing pad at *Rogi2* [31] (see Methods). We find that across integration methods, relative promoter strengths are comparable to those in transfection (Figures S2D to S2F). While absolute expression levels are reduced for low-copy lentivirus and landing pad integration, weak promoters exhibit higher mRNA and protein expression than expected in PiggyBac integration. This pattern may result from selection pressure from the co-expressed selection marker in the PiggyBac vector.

For many applications that require temporal control over expression, inducible promoters offer the advantage of small-molecule regulation of transcription. We characterized three promoters activated by small molecule-inducible transcriptional activators: doxycycline-inducible rtTA (Tet-On, CMV minimal promoter), rapamycin-inducible zinc-finger activator (COMET [24], YB TATA minimal promoter), and grazoprevir-inducible zinc-finger activator (synZiFTR [25], YB TATA minimal promoter). To facilitate comparison, each inducible promoter drives expression of mRuby2 with a bGH PAS. The cognate transcriptional activators for each synthetic promoter are expressed separately via the EFS promoter. We added small-molecule inducers at one day post-transfection and measured expression profiles via HCR Flow-FISH two days later (Figure 2E). Under these conditions, all three promoters display comparable levels of leaky expression in the absence of the inducer, similar to levels of expression from the weakest constitutive promoter, hPGK (Figure 2F). Interestingly, the induced Tet-On promoter has a similar expression level and effective translation rate to the constitutive CMV promoter. Both of these promoters contain the minimal CMV promoter, suggesting that this sequence drives both transcriptional and post-transcriptional kinetics. Upon induction, the Tet-On and synZiFTR promoters show modest increases in mRNA expression; however, the Tet-On promoter leads to a much higher level of protein expression (Figure 2F). This larger increase in protein level relative to mRNA level for the Tet-On promoter appears as a higher slope (Figure 2G) and effective translation rate (Figure 2H) compared to the other inducible promoters. Altogether, our data suggest that the differences in protein levels between promoters are impacted by differences in the 5’ UTR sequences in our constructs, which may impact translational dynamics independent of steady-state mRNA levels.

### Choice of polyadenylation signal impacts effective translation rate of mRNA transcripts

Given that 3’ UTR sequences can significantly impact RNA stability, protein translation, and lentivirus production efficiency, we sought to quantify the effects of polyadenylation signal (PAS) choice on expression kinetics [32–35]. We paired each constitutive promoter with three commonly used 3’ UTR sequences—bGH, derived from the bovine growth hormone gene; SV40, derived from the SV40 virus; and WPRE, derived from the woodchuck hepatitis virus (Figure 3A)—and transfected these transgenes into HEK293T cells. The bGH and SV40 PASs generate similar levels of protein (Figure 3C) and mRNA (Figure 3B). However, the WPRE sequence, which is not a mammalian PAS but enables the most efficient lentivirus production (Figure S2C), causes a reduction in protein levels for strong promoters (CAG and EF1α, Figure 3C) and an increase in mRNA levels for weak promoters (EFS and hPGK, Figure 3B).

**Figure 3.**
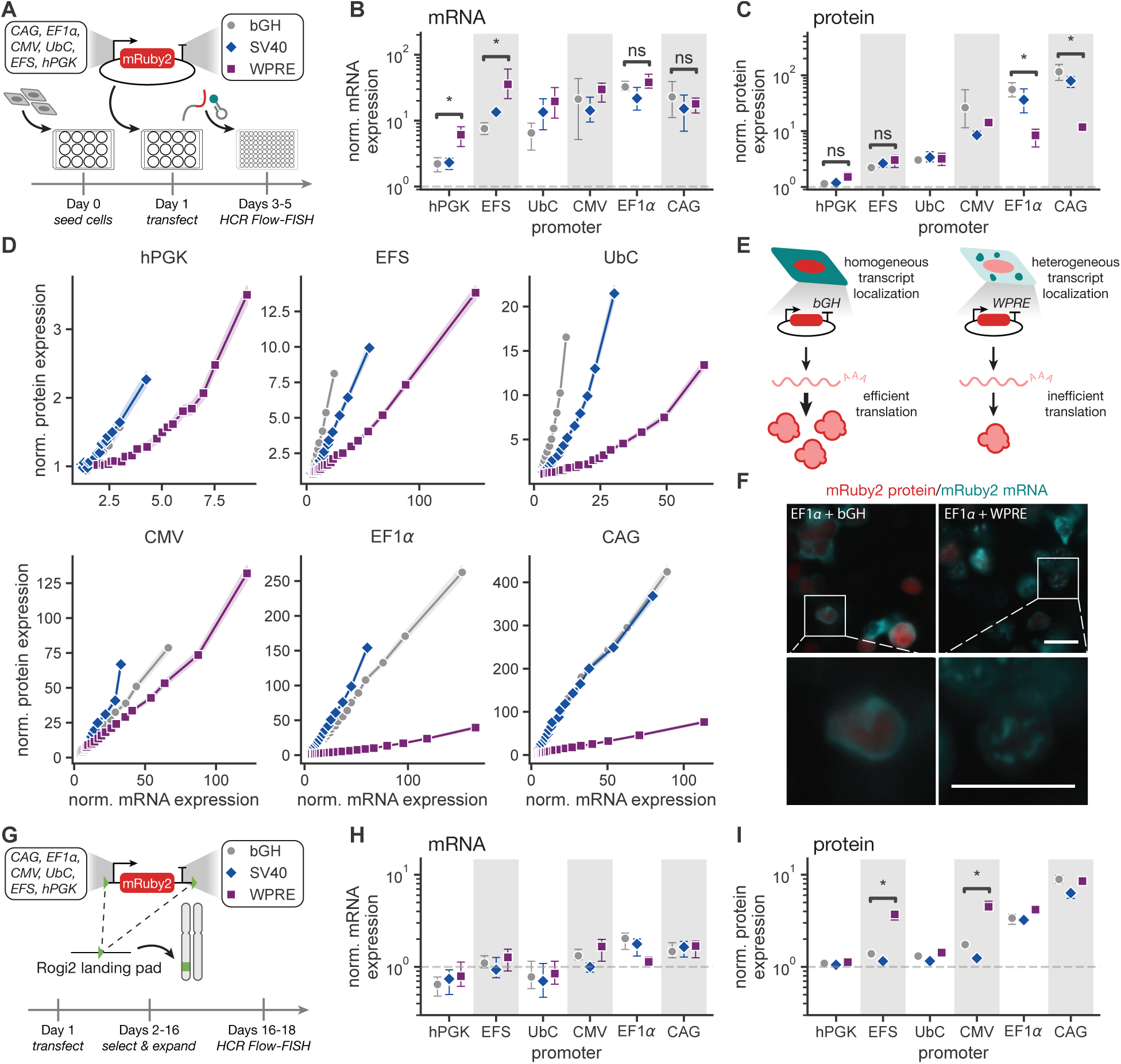
Choice of polyadenylation signal impacts effective translation rate of mRNA transcripts. **A.** HEK293T cells were transfected with a plasmid encoding mRuby2 driven by one of six different constitutive promoters along with one of three different 3’ UTR sequences. **B**, **C**. Normalized geometric mean of mRNA (Figure 3B) and protein (Figure 3C) fluorescence for varying promoter and PAS or 3’ UTR sequence in transfection of HEK293T cells. *: *p* ≤ 0.05, one-sided Mann Whitney test **D.** Normalized protein expression as a function of normalized mRNA expression for each promoter and 3’UTR pair. Data for one representative biological replicate is binned by marker level into 20 equal-quantile groups. Points represent geometric means of protein and mRNA expression for cells in each bin, and shaded regions represent 95% confidence intervals. **E.** Schematic of HCR RNA-FISH imaging of transfected HEK293T cells. Transcripts with a bGH PAS are distributed homogeneously throughout the cytoplasm and are efficiently translated. Transcripts with a WPRE sequence localize in puncta and produce less protein. **F.** HCR FISH imaging for HEK293T cells transfected and labeled with Alexa Fluor™ 514 HCR amplifiers. Scale bars represent 25 µm. Bottom row of images are the indicated zoomed in regions of the top images. **G.** The genes shown in Figure 3A were integrated site-specifically at a Rogi2 landing pad in HEK293T cells using Bxb1-mediated recombination. **H**, **I**. Normalized geometric mean of mRNA (Figure 3H) and protein (Figure 3I) fluorescence for varying promoter and PAS integrated at the Rogi2 locus in HEK293T cells. *: *p* ≤ 0.05, one-sided Mann Whitney test Normalized expression is calculated as the fold change of fluorescence intensity relative to a non-transfected sample. Points represent means of three biological replicates, and error bars represent the 95% confidence interval. All data is in arbitrary units from a flow cytometer.

Analyzing the data binned by marker expression, we find that the WPRE sequence results in the lowest slope of protein expression with respect to mRNA expression for all promoters, regardless of promoter strength (Figure 3D). This represents a lower effective translation rate for WPRE transcripts compared to bGH transcripts (Figure S3A). Therefore, the WPRE sequence leads to greater accumulation of mRNA transcripts, but these transcripts are translated at a lower rate. For strong promoters such as CAG and EF1α, which are already approaching the maximum transcriptional output in transfected cells, the decrease in effective translation rate results in a decrease in protein levels. For weak promoters such as EFS and hPGK, the increase in mRNA levels is balanced by the decrease in translation, resulting in relatively stable protein levels. With HCR FISH imaging, we observe that transcripts encoding WPRE form puncta, whereas bGH transcripts are distributed uniformly throughout the cytoplasm (Figures 3E and 3F). The presence of these puncta may indicate aberrant localization of WPRE transcripts, which may limit their rate of translation.

To examine how choice of 3’ UTR sequences affects expression profiles of integrated transgenes, we site-specifically integrated these cassettes into HEK293T cells at the *Rogi2* landing pad (Figure 3G). In contrast to the transfection results, we found that, at *Rogi2*, the WPRE sequence significantly increases protein levels for the CMV and EFS promoters despite minimal effects at the mRNA level (Figures 3H and 3I). However, the HCR Flow-FISH signal approaches the limit of detection, even for the strongest promoters. Therefore, we may not be able to resolve differences between mRNA levels at this low copy number using HCR Flow-FISH. Nevertheless, PAS choice significantly impacts the levels of protein expression from this promoter set at low copy number. Together, these findings demonstrate that PAS and 3’ UTR sequences tune mRNA and protein expression with different effects in transfection and integration.

### Gene coding sequence impacts effective translation rate but not mRNA levels

The identity of the transgene affects mRNA and protein levels via its specific sequence, which influences transcript stability, translation initiation, and protein stability. In particular, coding sequences may impact the secondary structure of the 5’ UTR. To investigate how the identity of the target gene affects expression profiles, we exchanged mRuby2 for tagBFP. While the mRuby2 and tagBFP transgenes are of similar length (711 and 702 bp, respectively), tagBFP has a modestly higher GC-content (56% versus 49% for mRuby2), which may result in differential mRNA nuclear export [36], stability [37], and translation efficiency [38]. Using our constitutive promoter panel to express tagBFP with a bGH PAS, we characterized the profiles of mRNA and protein expression in transfection of HEK293T cells (Figures S4 and 4A to 4C). While the slopes of the weak promoters show similar effective translation rates for tagBFP and mRuby2 (Figures 2C and 4B), tagBFP transcripts exhibit a different ordering of slopes for the set of strong promoters (Figures 2D and 4C).

**Figure 4.**
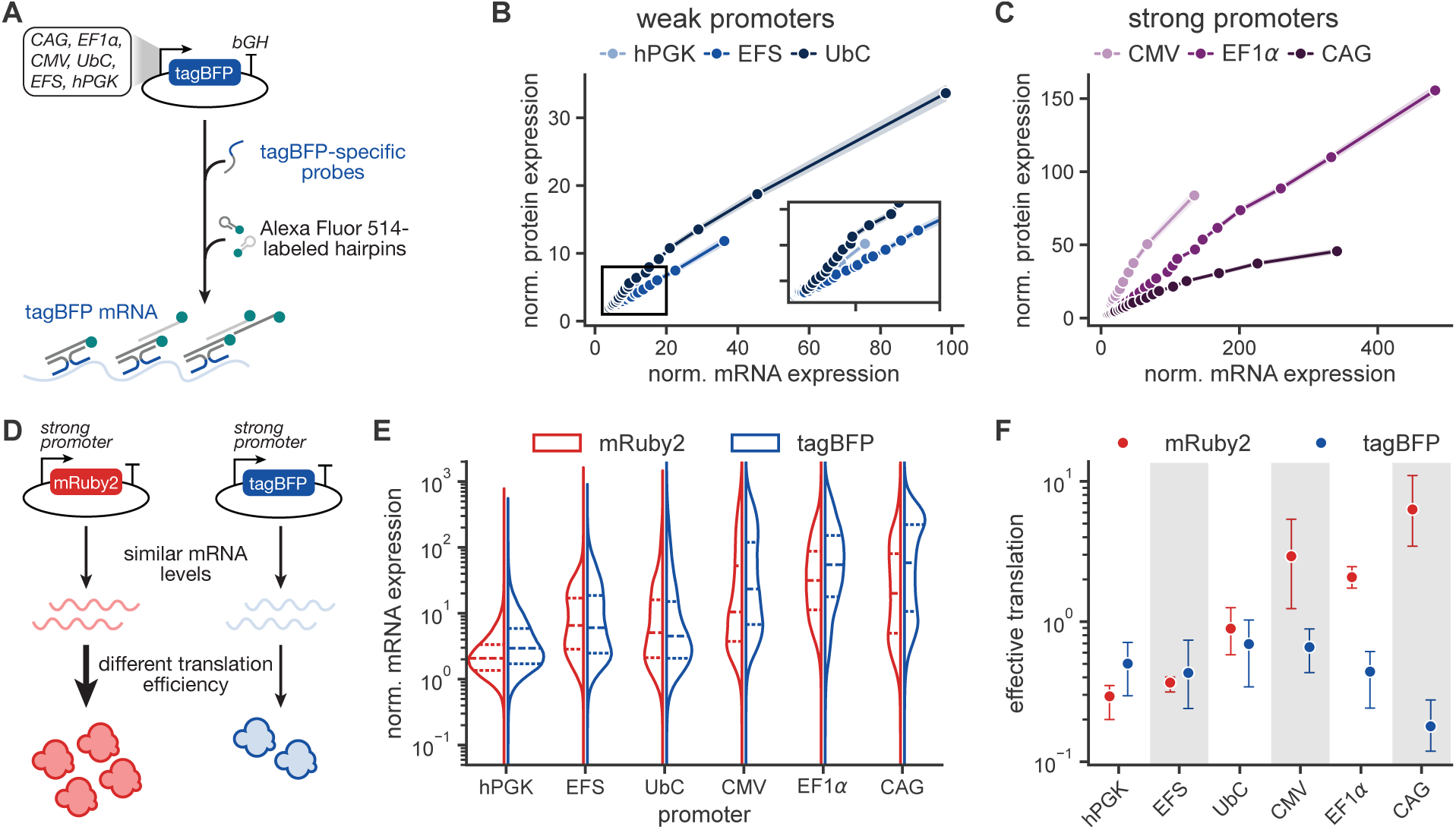
Gene coding sequence impacts effective translation rate but not mRNA levels. **A.** HEK293T cells were transfected with a plasmid encoding tagBFP driven by one of six different constitutive promoters (CAG, EF1_α_, CMV, UbC, EFS, or hPGK). **B.** Normalized protein expression as a function of normalized mRNA expression with three weak constitutive promoters. Inset axes show the indicated low expression domain. Data for one representative biological replicate is binned by marker level into 20 equal-quantile groups. Points represent geometric means of mRNA and protein levels for each bins. Shaded regions represent the 95% confidence interval. **C.** Normalized protein expression as a function of normalized mRNA expression for three strong constitutive promoters for one representative biological replicate. **D.** For genes with similar mRNA levels, differences in protein levels can indicate sequence-specific effects on the effective translation rate of RNA transcripts. **E.** Normalized expression distributions for mRuby2 and tagBFP mRNA (Alexa Fluor™ 514) with six constitutive promoters as measured by flow cytometry. **F.** Effective translation rate as calculated by the slope of a line fitted to the binned data. Normalized expression is calculated as the fold change of fluorescence intensity relative to a non-transfected sample. Points represent means of three biological replicates, and error bars represent the 95% confidence interval. All data is in arbitrary units from a flow cytometer.

To understand the differences between mRuby2 and tagBFP expression, we directly compared their RNA distributions (Figures 4A and 4E). Since both transcripts are labeled using the same HCR Flow-FISH amplifiers, we can quantitatively compare mRNA profiles generated by HCR Flow-FISH. Strikingly, each constitutive promoter generates similar mean RNA levels regardless of the transgene expressed (Figure 4E). Similar to mRuby2, tagBFP RNA levels show more variance than tagBFP protein levels (Figure S4). Thus, we find substitution of tagBFP for mRuby2 does not substantially affect the profiles of mRNA.

While the levels of mRNA are comparable between coding sequences for each promoter, protein levels differ substantially across the stronger promoters (Figure S4B). The strongest promoters (EF1α and CAG) display lower levels of tagBFP protein relative to CMV. Lower expression of tagBFP protein cannot be explained by potential differences in protein half-life because a reduction in protein half-life would result in lower protein expression across all promoters. Rather, these data suggest that the promoters affect the translation of tagBFP transcripts (Figure 4D). Curiously, unlike mRuby2, we observe that tagBFP exhibits a lower effective translation rate for strong promoters than for weak promoters (Figure 4F). The consistency in mRNA levels between mRuby2 and tagBFP suggests that gene identity has minimal effect on transcription. Rather, we suggest that the sequences of these coding regions may impact RNA processing, transcript localization, RNA stability, or translation that manifest as differences in protein levels. Further characterization of 5’ UTR architectures may allow for the identification of factors impacting translation [39].

### Examining transcript isoforms highlights promoter-specific patterns of gene regulation

The UTRs of transcripts can have sequence-specific effects on mRNA transport, translation and stability [40]. Additionally, introns within the 5’ UTRs associated with synthetic promoters, such as CAG, EF1α, and UbC, may affect transcription and mRNA processing kinetics via “intron-mediated enhancement” [41]. We sought to define these effects by precisely mapping transcript start sites (TSS) and transcription end sites (TES), as well as splice site positions, using long-read direct RNA sequencing (Figure 5A). We sequenced transcripts from six cell lines randomly integrated with transgenes containing different constitutive promoters, which we previously analyzed with HCR Flow-FISH and RT-qPCR (Figures 1F, S2D and 5B). We find that transcript isoforms are highly uniform. For the set of constitutive promoters, only a small fraction of transcripts deviate from expected transcript start and end positions (Figures S5A and 5C).

**Figure 5.**
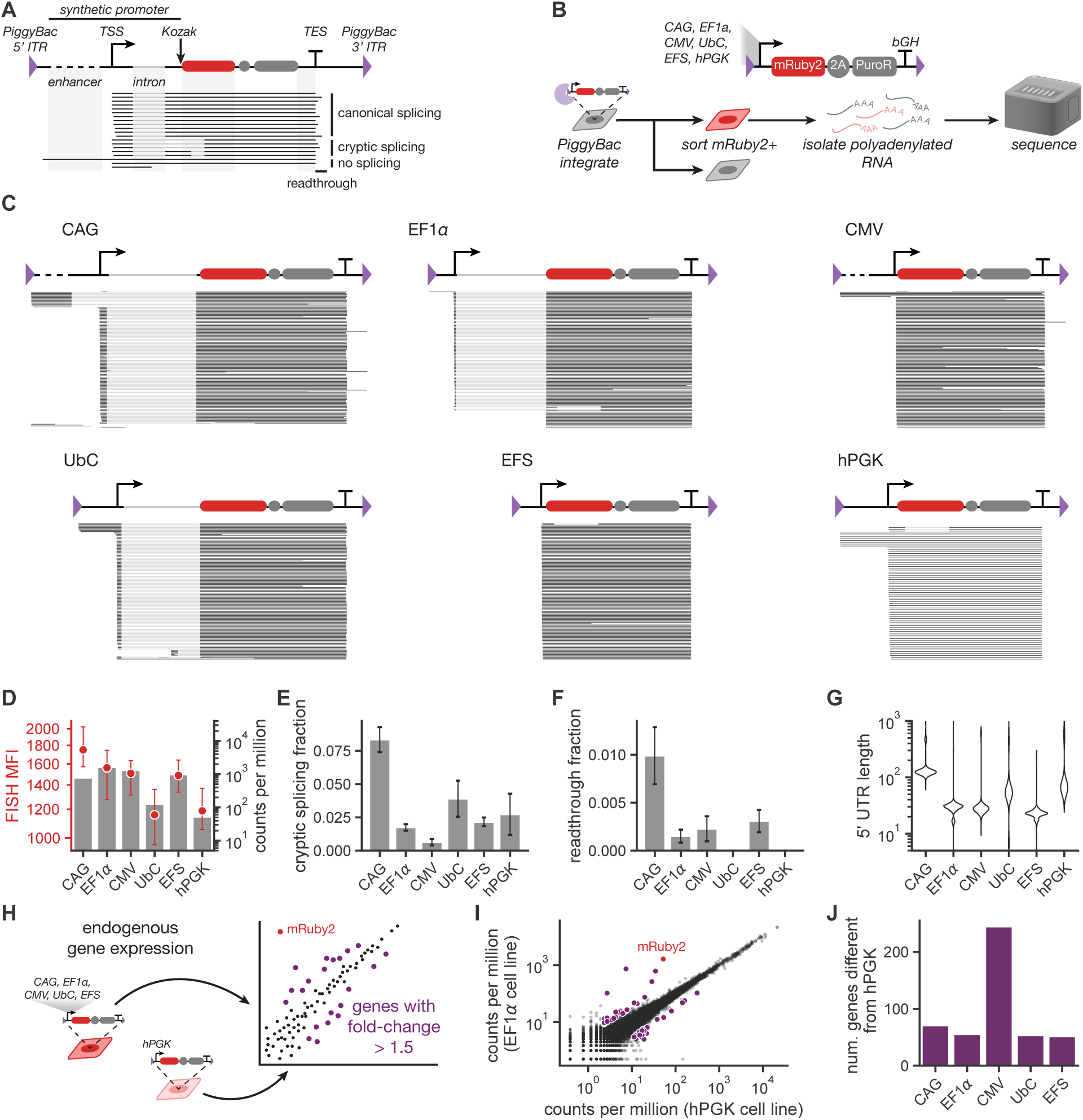
Examining transcript isoforms highlights promoter-specific patterns of gene regulation. **A.** Example transcripts depicting a synthetic promoter sequence containing an intron and enhancer. Each line represents a separate sequencing read. The TSS and TES are the locations where the majority of reads begin and stop, respectively. Reads are separated by splicing status, where canonical splicing refers to reads with the annotated intron in the promoter region and cryptic splicing refers to reads using a cryptic intron in the coding region. Readthrough transcripts extend past the annotated TES. **B.** Genetic cargoes were integrated randomly into HEK293T cells using PiggyBac transposase. Cells were sorted to yield polyclonal, mRuby2+ cell lines. RNA was isolated from each cell line and subjected to long-read sequencing. **C.** RNA transcript maps for the six constitutive promoters tested. All samples were downsampled to display 144 reads for consistency, aside from hPGK, for which there are only 77 total reads. All reads are shown in Figure S5A. **D.** HCR Flow-FISH MFI (left axis, points) and mRuby2 sequencing counts per million (right axis, bars) for the six cell lines. MFI is displayed in arbitrary units as the mean of three biological replicates ± the 95% confidence interval. **E**, **F**. Fraction of reads exhibiting cryptic splicing (**E**) or readthrough (**F**). Error bars represent the standard deviation assuming counts follow a binomial distribution. **G.** Distribution of 5’ UTR lengths across reads. **H.** Endogenous genes are considered “differentially expressed” between the cell lines if the absolute value of the fold change is greater than 1.5. These genes are indicated in purple. mRuby2 expression is indicated in red. **I.** Endogenous gene expression levels for cell lines integrated with the EF1_α_ (y-axis) or hPGK promoters (x-axis). Data for the rest of the promoters is shown in Figure S5B. Lists of these genes are reported in Table S2. **J.** Number of differentially expressed endogenous genes in each engineered cell line relative to hPGK.

Despite the possibility of substantial differences in local sequence, gene, or chromatin context due to random integration into the genome, we observe very little evidence for transcriptional read-through from an upstream promoter or for transcriptional differences caused by local effects. Additionally, transcript counts from sequencing agree with previous measures of mRuby2 mRNA levels via HCR Flow-FISH (Figure 5D). A very small fraction of reads exhibit unexpected splicing patterns, where aberrant splicing results in the use of a cryptic splice site in the mRuby2 coding sequence (Figure 5E). An even smaller fraction of reads display readthrough past the expected transcription end site. However since direct RNA sequencing only captures polyadenylated molecules, we lack the ability to estimate any readthrough transcripts that are not polyadenylated.(Figure 5F). The distribution of 5’ UTR lengths differs across the set of constitutive promoters (Figure 5G).

We defined the 5’ UTR as the distance between the observed TSS and the mRuby2 start codon. Specifically, CAG, UbC, and hPGK have the longest 5’ UTRs at ∼50-100 nucleotides, while the 5’ UTRs of EF1α, CMV, and EFS are ∼20-30 nucleotides. Of the promoters tested, UbC and hPGK had the lowest number of mRuby2 reads (Figure 5D). Potentially, the longer 5’ UTRs associated with these promoters may reduce transcript stability as has been observed natively [42].

Finally, we evaluated the relative impact of each promoter on endogenous gene expression by comparing to gene expression in the engineered cell line with the hPGK cassette (see Methods). hPGK is the weakest constitutive promoter, which we would expect to exert the lowest impact on endogenous expression (Figure 5H). Overall, expression of endogenous genes is highly correlated across cell lines. Comparing to the integrated line with the transgene driven by hPGK, the correlation indicates that transgenes exert minimal impact on native genes (Figures S5B, 5I and 5J and Table S2). Notably, CMV induces the most variability in endogenous gene expression, affecting genes associated with viral response pathways (Table S2). Overall, these results indicate that these promoters may exert minimal burden while producing a range of expression levels in HEK293T cells.

## Discussion

Properly tuning transgene expression levels is essential to synthetic circuit design. Selecting a promoter—commonly, either a strong native promoter, a viral-derived promoter, or a synthetic promoter that recruits transcriptional activator proteins [24, 25, 43]—is often the method of choice to change the rate of transcription. However, promoter choice can be a blunt tool, as mRNA processing, mRNA transport, translational initiation, and protein stability can all contribute to levels of transgene expression (Figure 1). In this work, we simultaneously measured mRNA and protein levels in order to assess impacts on both transcription and translation at steady state. Indeed, we find that strong promoters both transcribe more mRNA and have higher effective translation rates (Figure 2). To explore how other experimentally-accessible, genomically-encoded variables affect expression, we investigated how the polyadenylation signal in the 3’ UTR (Figure 3), coding sequence identity (Figure 4), and 5’ UTR identity and length (Figure 5) jointly determine transgene levels.

In prokaryotes, steady-state mRNA transcript levels strongly correlate with translation initiation rate, though RNA secondary structures such as hairpins, G-quadruplexes, and i-motifs in the 5’ and 3’ UTRs all affect translation [44]. In eukaryotes, the transcriptional landscape is more complicated, with processes such as splicing, polymerase termination, polyadenylation, and transcription degradation driven from relatively opaque sequences with no discernible secondary structure. In fact, the sequence of the 5’ UTR encoded by a synthetic promoter may impact both the translation rate and mRNA stability, of the transgenic transcripts in human cells [40, 45, 46].

Within our panel of promoters, splicing within the 5’ UTR increases effective translation rates, consistent with previous observations of splicing-mediated enhancement of gene expression [30, 41, 47–49]. Specifically, the effective translation rate of EF1α is six times higher than that of EFS, which has the EF1α intron removed (Figure 2). As confirmed by long-read sequencing, the post-spliced 5’ UTRs of transcripts expressed from these promoters differ by only 10 nucleotides. This difference may result from the recruitment of splicing factors that aid in not only splicing but also nuclear export of the mRNA transcripts, as has been observed with native genes [41, 47].

A growing number of synthetic circuit designs place synthetic target sites for post-transcriptional control in the 5’ or 3’ UTR or utilize synthetic introns to encode for circuit components [7, 10, 50]. Understanding how the transcript isoform influences transcript processing and translation will inform design of robust circuits. Here, we find that mature RNA isoforms are highly uniform (Figure 5). Remarkably, only a small fraction of mature transcripts have non-canonical TSS or TES usage, splicing patterns, or readthrough, suggesting that it is reasonable to conceptually model each gene as its dominant isoform, instead of having to rely on a more complicated population model that accounts for isoform diversity. Rather, variability in transcription and translation rates may explain the variation in RNA and protein levels within a design. Since direct RNA sequencing quantifies only mature transcripts—those that are polyadenylated—we cannot assess the extent of immature, unprocessed transcripts that are produced. However, given the inherent instability of transcripts lacking a polyA tail, these transcripts are likely quickly degraded and would not contribute substantially to steady-state mRNA or protein levels.

As polyadenylation signal putatively affects mRNA stability, we investigated three commonly used 3’ UTR sequences used in synthetic biology; two viral-derived sequences (WPRE, SV40) and one native sequence (bGH). All three are commonly used and are often arbitrarily paired to reduce the probability of plasmid recombination. While the SV40 and bGH PASs behave similarly,(Figures 3B and 3D), we find that WPRE, a virally-derived sequence that is necessary for lentiviral production (Figure S2), localizes mRNA transcripts to puncta in the cytoplasm and decreases the effective translation rate (Figure 3). Potentially, these puncta may represent specific subcellular structures such as stress granules or P bodies [51, 52]. For gene circuits that act at the post-transcriptional level (such as a microRNA-based iFFL [10] or an antisense integral controller [8]), the localization of mRNA transcripts may limit performance by altering the local concentrations of circuit components. Further study of mRNA localization can better inform selection of 3’ UTR elements to maximize circuit response and minimize variability. Nevertheless, we observed that these PAS effects may depend on copy number or integration context (Figure 3).

When comparing the expression profiles of promoters between transfection and integration systems, we identified cases where the transfection profile offers reasonable predictability of expression within the integrated context. While complex coupling can occur on multi-gene plasmids [53], single-gene plasmids are free from coupling to adjacent genes and may offer a rapid prototyping platform for transcriptional units. Particularly with the same pairings of promoter and PAS, we found that transfection experiments could predict the relative expression profiles for integrated transgenes, albeit with different absolute levels of expression (Figure S2). Interestingly, we found that weak promoters—particularly EFS—exhibit higher mRNA and protein expression in PiggyBac integration than predicted by transfection (Figure S2D). The selection pressure applied during cell line creation may cause the expression from weaker promoters to skew higher due to the cotranscriptional expression of an antibiotic resistance gene. Taking into account the time-intensive workflow for cell line generation and the limitations of HCR Flow-FISH detection at low mRNA copy copy number, characterization in transfection offers a way to quickly assess mRNA distributions with higher resolution than in low-copy number cell lines. Therefore, transfection characterization results have predictive power as long as promoter, gene, and PAS sequences remain the same (Figures S2 and 4). This predictability is important for the development of gene circuits that function reliably in engineered cell lines.

Our study also allowed us to consider how transgene expression interacts with endogenous gene expression in engineered cell lines. First, we observed very consistent transcription start and end site usage across transgenes randomly integrated in the genome (Figure 5C), demonstrating that the local sequence context does not heavily impact transgene expression through readthrough from upstream promoters and/or genes. Second, we find that there is very little variability in endogenous gene expression across cells with integrated transgenes driven by different promoters of different strengths, suggesting that transgene expression does not cause competition for cellular resources (Figure 5 and Table S2). Together, our two findings suggest that transgenes are often self-contained gene regulatory units that exert minimal impact on native gene regulatory mechanisms and networks. Further investigation of the durability of transgene expression over time and characterization of effects on endogenous genes in more cell types could aid in identifying genetic elements with reliable performance in different contexts.

With the increasing number of studies using library screening to identify genetic elements [11, 39, 54], HCR Flow-FISH might be adapted to screen larger libraries of genetic sequences for desired profiles of RNA expression. However, while we have demonstrated the value of high-resolution profiling with HCR Flow-FISH and direct RNA sequencing via long-reads for a handful of transgenes, scaling these methods to hundreds or thousands of designs remains unfeasible. In its current embodiment, HCR Flow-FISH characterization only requires basic biochemistry and access to a flow cytometer. However, the handling time over a three-day workflow limits throughput to ∼100 transgene variants. Integration of HCR Flow-FISH with a microfluidic platform with automated wash steps would greatly increase capacity and could reduce variability in mRNA labeling, supporting broader adoption into characterization workflows [26].

Through single-cell profiling of RNA and protein expression as well as analysis of RNA isoforms, we identify determinants of transgene expression levels in engineered cells and interactions of these transgenes with endogenous gene expression. We find that promoter choice for transgene expression influences both RNA transcript abundance and effective translation rate. The effective translation rate is further affected by the transgene coding sequence and 3’ UTR sequence. Differences in effective translation rates from constructs may be larger across cell types and show cell-type specific profiles of expression, potentially explaining the heuristic choice. Our framework of profiling RNA and protein levels simultaneously in single cells can be expanded to additional cell types and genetic elements to identify new sequences as well as tune expression for diverse functions. Increasing knowledge of how genetic elements contribute to profiles of expression will support predictive design of programmable gene circuits with controlled functions in diverse cell types.

## Acknowledgments

Research reported in this manuscript was supported by the National Institute of General Medical Sciences of the National Institutes of Health under award numbers R35-GM143033 and R35-GM133762, by the National Science Foundation with the NSF-CAREER under award numbers 2339986 and 2237568, by the Institute for Collaborative Biotechnologies, and by the Air Force Research Laboratory MURI under award number FA9550-22-1-03t16. E.L.P, K.S.L, and S.R.K. are supported by the National Science Foundation Graduate Research Fellowship Program under grant number 1745302. Work completed at the MIT BioMicro Center was supported in part by the Koch Institute Support (core) Grant P30-CA014051 from the National Cancer Institute.

The authors thank Adam Beitz, Nat Wang, Brittany Lende-Dorn, Mary Ehmann, Jane Atkinson, and Maria Castellanos for their helpful feedback on the manuscript.

## Author Contributions

K.E.G., E.L.P., D.S.P., K.S.L., C.P.J., and S.R.K. conceived and outlined the project. C.G.O. performed initial HCR Flow-FISH validation. E.L.P., D.R.G., and K.S.L. performed HCR Flow-FISH quantification. E.L.P. and D.R.G. performed HCR RNA-FISH imaging. D.S.P. performed modRNA synthesis, Rogi2 LP cell line creation, and site-specific integration. E.L.P. and D.R.G. performed transfection of constitutive promoters. K.S.L. performed transfection of inducible promoters. E.L.P. and C.P.J. performed PiggyBac integration and sorting. C.P.J. and S.R.K. performed lentivirus production and transduction. V.S. and R.F.D. generated and analyzed long-read sequencing data. E.L.P., K.S.L., and C.P.J. performed data analysis. E.L.P. wrote the initial draft. All authors wrote relevant methods, edited the draft, and helped in manuscript preparation. K.E.G. and A.A.P. supervised the project.

## Materials and Methods

### Cloning

Expression plasmids were generated using a multi-level Golden Gate cloning scheme. First, individual genetic part fragments were amplified or digested from commercial DNA sources. Each fragment was inserted into a corresponding part positioning vector (pPV) backbone via Gibson Assembly using Hifi DNA Assembly Master Mix (NEB, M5520). On each pPV, the region of interest is flanked by BsaI restriction sites detailed in Table S3. A full list of pPVs used in this work and their DNA sources are reported in Table S4. Full transcriptional units were assembled in a BsaI (NEB, R3733L) Golden Gate reaction to yield the kanamycin-resistant plasmids (pShips) used in transfection experiments. In a second round of PaqCI (NEB, R0745L) Golden Gate reactions, full transcriptional units were inserted into backbones for genomic integration (pHarbors) Bxb1 recombinase, PiggyBac transposase, and lentivirus.

To clone the homology-directed repair (HDR) donor template for targeting Rogi2 with a Bxb1 landing pad (LP), 5’ and 3’ Rogi2 homology arms were PCR amplified from HEK293T genomic DNA. To facilitate PCR genotyping of CRISPR-edited clones, the length of the homology arms were selected based on the co-design of randomly-generated 5’ and 3’ barcode sequences and genotyping primers with primer pairs that (1) flank the HDR junction of the donor DNA and the Rogi2 locus and (2) were not predicted by PrimerBLAST to produce off-target amplicons of similar size. The resulting barcodes were encoded on the primers used to amplify the Rogi2 5’ and 3’ homology arms. These were assembled with pHarbor backbone fragments PCR amplified with primers that encoded the Rogi2 protospacer and protospacer adjacent motif (PAM) sequences. Fragments were designed such that each homology arm is flanked by the Rogi2 protospacer and PAM sequence needed for In Trans Paired Nicking (ITPN) editing [55]. The PCR fragments were assembled into a pHarbor using an Esp3I (NEB, R0734L) Golden Gate reaction.

The LP architecture inserted at Rogi2 was based on the STRAIGHT-IN platform [56] with modifications. The LP consists of two transcriptional units Figure S6. The 5’ unit consists of an EF1α-BsdR-bGH cassette, which confers blasticidin resistance and is used for selecting cells which underwent HDR. The 3’ unit consists of a Bxb1 attB site and a PuroR-bGH cassette lacking a promoter and start codon, which is used for enriching Bxb1-mediated integration of attP donor plasmids in the LP cell line. Each transcriptional unit was assembled in a BsaI Golden Gate reaction to generate pShips, which were subsequently assembled into the Rogi2-targeting pHarbor in a PaqCI Golden Gate reaction, yielding the final ITPN donor plasmid used to create the HEK293T Rogi2 LP cell line.

### Cell culture

#### HEK293T

HEK293T cells (ATCC, CRL-3216) were cultured using DMEM (Genesee Scientific, 25-501) plus 10% FBS (Genesee Scientific, 25-514H) and incubated at 37°C with 5% CO2. Cells were passaged at 80-90% confluence, in which spent media was aspirated, and cells were washed with PBS (Sigma-Aldrich, P4417-100TAB) then subsequently dissociated with 0.25% Trypsin-EDTA (Genesee Scientific, 25-510) diluted in PBS. After four minutes, cells were spun down at 400 rcf for 5 minutes, resuspended in media, then transferred to a new flask. Media was replaced with fresh DMEM + 10% FBS every 2-3 days.

#### CHO-K1

CHO-K1 cells (ATCC, CCL-61) were cultured using DMEM/F12 (Corning, 10-090-CV) plus 10% FBS and incubated at 37°C with 5% CO2. CHO-K1 cells were passaged identically to HEK293T cells.

#### iPS11

iPS11 cells (Alstem, iPS11) were cultured using mTeSR™ Plus (STEMCELL Technologies, 100-1130) on Geltrex™-coated (ThermoFisher Scientific, A1413302) plates and incubated at 37°C with 5% CO2. Cells were passaged in clumps using ReLeSR™ (STEMCELL Technologies, 100-0484) according to the manufacturer’s instructions.

### Transfection

#### HEK293T

In preparation for experiments, HEK293T cells were counted using a hemocytometer, seeded with 0.1% gelatin coating (Sigma-Aldrich, G1890-100G) at a density of 150,000 cells per 12-well or 25,000 cells per 96-well, and transfected 24 hours later. For imaging experiments, cells were seeded in Geltrex-coated glass-bottom 96-well plates. Transfection was performed using linear polyethylenimine, PEI (Fisher Scientific, 4389603). Transfection mixes were prepared using a ratio of 4 µg PEI to 1 µg DNA. First, a master mix of PEI and KnockOut™ DMEM (ThermoFisher Scientific, 10-829-018) was prepared and incubated for a minimum of 10 minutes. This master mix was then added to DNA mixes containing 450 ng of output plasmid and 450 ng of transfection marker plasmid per 12-well (or 56 ng of each plasmid per 96-well). These conditions mixes were further incubated for 10 to 15 minutes and then added on top of the growth media in the plate.

For experiments with constitutive promoters, media was replaced with fresh DMEM + 10% FBS after 24 hours. At 2 days post transfection, HEK293T cells were dissociated by adding 0.25% Trypsin-EDTA (diluted in PBS) for four minutes, followed by quenching with an equal volume of DMEM + 10% FBS. After centrifuging at 500 rcf for 5 minutes, cells were resuspended in PBS and transferred to a v-bottom plate for flow cytometry or subsequent HCR Flow-FISH.

For experiments with inducible promoters, 24 hours after transfection media was replaced with fresh DMEM + 10% FBS containing the corresponding small molecule inducer or solvent control. Inducer stocks were prepared as follows: doxycycline (dox; Sigma-Aldrich, D3447) in water at 1 mg/mL, grazoprevir (GZV; MedChem Express, HY-15298) in DMSO at 1 mM, and rapamycin (Rap; Millipore Sigma, 553210) in DMSO at 200 µM. Grazoprevir and rapamycin stocks were stored at −80°C until use, then kept at 4°C for up to two weeks. Doxycycline stocks were stored at −20°C until use, then kept at 4°C. Small molecule stocks were diluted in DMEM + 10% FBS to the following concentrations for experiments: 1 µg/mL doxycycline, 1 µM grazoprevir, and 0.1 µM rapamycin. Two days after small molecule addition (3 days post-transfection), HEK293T cells were dissociated by adding 0.25% Trypsin-EDTA (diluted in PBS) for four minutes, followed by quenching with an equal volume of DMEM + 10% FBS. After centrifuging at 500 rcf for 5 minutes, cells were resuspended in PBS and transferred to a v-bottom plate for flow cytometry or subsequent HCR Flow-FISH.

#### CHO-K1

Similar to HEK293T cells, CHO-K1 cells were seeded 24 hours prior to transfection with 0.1% gelatin coating at a density of 25,000 cells per 96-well. Using PEI, 56 ng of output plasmid and 56 ng of transfection marker plasmid were delivered to each 96-well. Media was replaced with fresh DMEM/F12 + 10% FBS after 24 hours. At 2 days post transfection, CHO-K1 cells were dissociated, resuspended in PBS, and transferred to a v-bottom plate for flow cytometry.

#### iPS11

For transfection experiments, iPS11 cells were dissociated using Gentle Cell Dissociation Reagent (STEMCELL Technologies, 100-1077) according to manufacturer’s instructions and counted using a hemocytometer. Cells were plated 48 hours prior to transfection in mTeSR™ Plus with 10 µM ROCK inhibitor (Millipore Signma, Y0503-5MG) and 100 U/mL penicillin-streptomycin (Gibco, 15140122) at a density of 15,000 cells per 96-well. After 24 hours, ROCK inhibitor was removed. On the day of transfection, transfection mixes were prepared with FUGENE® HD (FuGENE, HD-1000) using a ratio of 3 µL reagent to 1 µg DNA, and the media was changed to Opti-MEM™ (ThermoFisher Scientific, 31985062). 50 ng of output plasmid and 50 ng of transfection marker plasmid were delivered to each 96-well. Fresh mTeSR™ Plus with penicillin-streptomycin was added 4 hours after transfection, and the media was changed 24 hours after transfection. At 2 days post transfection, cells were dissociated using Gentle Cell Dissociation Reagent, resuspended in PBS, and transferred to a v-bottom plate for flow cytometry.

### modRNA synthesis and titration curve experiments

The mRuby2 modRNA used in this study was synthesized from a plasmid template which encoded the 5’-UTR of human β-globin, a Kozak sequence, the coding sequence for mRuby2, and the 3’-UTR of human β-globin. The linear template for *in vitro* transcription (IVT) was generated via PCR using Q5 DNA Polymerase (New England Biolabs, M0491) with the forward primer (5’-AGCTATAATACGACTCACTATAAGctcctgggcaacgtgctg-3’) encoding the T7 promoter (upper-case bases) and binding the 5’ UTR (lower-case bases) and the reverse primer (5’-poly(T)116-GCAATGAAAATAAATGTTTTTTATTAGGCAGAAT-3’) encoding the poly(A) tail and binding the 3’-UTR. The PCR product was isolated on a 1% agarose gel, excised, and purified using the Monarch PCR and DNA Cleanup Kit (New England Biolabs, T1030). 200 ng of purified product served as template in a 20 µL IVT reaction using the HiScribe T7 High Yield RNA Synthesis Kit (New England Biolabs, E2040), fully substituting UTP with N1-methylpseudouridine-5’-triphosphate (TriLink Biotechnologies, N-1081) and co-transcriptionally capping with CleanCap Reagent AG (TriLink Biotechnologies, N-7114). IVT reactions were incubated at 37°C for 4 hours, at which point reactions were diluted to 50 µL, treated with 2 µL DNase I (New England Biolabs, M0303), and incubated at 37°C for 30 min to degrade the IVT PCR template DNA. Synthesized modRNA was column purified and eluted with 60 µL water using the 50 µg Monarch RNA Cleanup Kit (New England Biolabs, T2040). A small sample was nanodropped and ran on a native denaturing gel to determine modRNA concentration and verify full-length product. The modRNA was dispensed in single-use aliquots and stored at −80°C.

For modRNA titration experiments, HEK293T cells were seeded on 12-well plates with 150,000 cells per well three days before modRNA transfection. Two days before modRNA transfection, plasmid control conditions were transfected as described above. One day before modRNA transfection, media was replaced with fresh DMEM + 10% FBS. The following day, each modRNA mixture was transfected in triplicate using 1.6 µL Lipofectamine MessengerMAX (Thermo Fisher Scientific, LMRNA) according to manufacturer’s instructions. Each well was transfected with varying amounts of mRuby2 modRNA. To normalize transfection efficiency across conditions, each condition was adjusted to a total modRNA amount of 800 ng with a Cre-encoding modRNA that has no predicted affinity with the mRuby2 FISH probes. Four hours after modRNA transfection, cells were dissociated for HCR Flow-FISH by adding 0.25% Trypsin-EDTA (diluted in PBS) for four minutes, followed by quenching with an equal volume of DMEM + 10% FBS.

### Site-specific integration

#### Generation of the HEK293T Rogi2 Bxb1 landing pad cell line

We generated a Bxb1 attP landing pad (LP) cell line for facile site-specific integration of genetic cargoes in HEK293T cells. We chose the *Rogi2* locus for integration as it is far from other genes and regulatory elements [31]. To perform the genomic integration, we leveraged In Trans Paired Nicking (ITPN) [55]. This CRISPR-based strategy flanks the Rogi2 homology arms on the donor DNA plasmid with the cognate Rogi2 protospacer sequence targeted on the genome, 5’-CATCAGACTTGATAGCACTG<u>AGG</u>-3’ (PAM underlined). Subsequent *in situ* nicking of the Rogi2 locus and the donor DNA plasmid by a high-fidelity Cas9 nickase variant facilitates precise installation of donor DNA at the target locus while minimizing random integration events associated with double-strand breaks generated by wild type Cas9.

To implement ITPN, we adopted a staggered delivery approach similar to CRISPR for long-fragment integration via pseudovirus (CLIP) [57], where the donor DNA is delivered first, followed by delivery of CRISPR components 24 hours later (Figure S6A). One day before transfection, 150,000 HEK293Ts were plated on a single 12-well. Cells were then transfected with 1000 ng of donor DNA plasmid using PEI. The next day, cells were transfected with 300 ng of nCas9(1.1) modRNA (synthesized in-house with the HiScribe T7 High Yield RNA Synthesis Kit, New England Biolabs, E2040) and 100 ng of Rogi2 sgRNA (synthesized with the EnGen sgRNA synthesis kit, New England Biolabs, E3322) using 0.8 µL of Lipofectamine MessengerMax.

Two days after RNA delivery, cells were passaged onto a single 6-well. The following day, cells were treated with 10 µg/mL blasticidin (Tocris, 5502) for four days to enrich cells which genomically integrated the donor DNA. Single cells from the polyclonal population were sorted into individual wells of a 96-well plate using the Sony MA-900 flow sorter. Two weeks post-sort, confluent monoclonal lines were passaged to 24-well plates, with half of the cells harvested for genotyping PCR. Harvested cells were pelleted and resuspended in 50 µL Cell Lysis Buffer (10X) (Cell Signaling Technology, 9803S) and 0.5 µL of Proteinase K (New England Biolabs, P8107S). Cells were lysed by incubating the suspension for 45 minutes at 85°C.

As amplifying the Rogi2 locus proved challenging, we developed an efficient PCR screening method to detect for the desired insertion at Rogi2. This consisted of designing the donor DNA with short barcode sequences located between the attP Landing Pad and the Rogi2 homology arms (Figure S6B). These barcode sequences were co-designed with the genotyping primers using PrimerDesign (NCBI) to minimize potential off-target amplicons. To enhance PCR sensitivity, we used nested PCR [58]. For the first, outer PCR, 1 µL of the cell lysate was used as template in a 20 µL PCR using Apex Taq RED Master Mix, 2X (Genesee Scientific, 42-138) with a 30 second extension time and a “touchdown” annealing temperature [59]. This consisted of setting the annealing temperature of the first PCR cycle at 72°C, with each subsequent PCR cycle decreasing the annealing temperature by 1°C until reaching a final annealing temperature of 57°C, followed by an additional 12 cycles at this annealing temperature. 1 µL of the first PCR was used as template for a second, inner PCR, following the same PCR conditions. PCR products obtained from the second PCR reaction were resolved on a 2% agarose gel (Figure S6C).

After identifying monoclones which passed the genotyping PCR screen, each candidate was phenotypically screened for the ability to effectively integrate and express genetic cargoes encoded on Bxb1 attB donor plasmids via Bxb1-mediated recombination. Clone #14 emerged as the best candidate from this screen, hereafter referred to as Rogi2 LP, and was subsequently used in this study.

#### Bxb1-mediated integration of cargoes in the HEK293T Rogi2 landing pad cell line

The procedure outlined here functions analogous to the STRAIGHT-IN iPSC landing pad platform (Figure S6D) [56]. The Rogi2 LP line contains a puromycin resistance gene missing a promoter and start codon. Upon Bxb1-mediated recombination between the attB site on the donor plasmid (Figure S6E) and attP site in the LP, an EF1α promoter and start codon is placed in-frame of the resistance gene, conferring recombined cells resistance to puromycin (Figure S6F). A total of 18 different donor plasmids were cloned encoding mRuby2 driven by six different promoters (CAG, EF1α, CMV, UbC, EFS, or hPGK) and three different PAS sequences (bGH, SV40, or WPRE). To integrate the donor plasmids, Rogi2 LP was plated on 24-well plate at a seeding density of 75,000 cells per well one day before transfection. Cells were then co-transfected with 300 ng of donor attB plasmid and 200 ng of CAG-Bxb1 (gift from the Wong Lab at Boston University) using PEI. Once confluent (two to three days post-transfection), cells were passaged onto a 6-well, with puromycin (1 µg/mL, Invivogen, ant-pr-1) administered the following day. Once confluent (five to six days post puromycin selection), cells were passaged at a split ratio of 1:10 to dilute out residual donor plasmid, at which point cells were ready for use in downstream analyses.

### PiggyBac integration

Six different genetic cargoes were randomly integrated into HEK293T cells, encoding expression of mRuby2-2A-PuroR-bGH driven by CAG, EF1α, CMV, UbC, EFS, or hPGK. For PiggyBac transposase-mediated integration, 100,000 HEK293T cells per well were seeded in a 24-well plate coated with 0.1% gelatin. Each well was transfected as described above with 225 ng of donor plasmid and 225 ng of PiggyBac transposase plasmid (gift from the Elowitz Lab). At one day post-transfection, media was replaced with fresh DMEM + 10% FBS. At two days post-transfection, cells in each 24-well were passaged to a 6-well. One day after passaging, media was replaced with fresh DMEM + 10% FBS with 1 µg/mL puromycin for selection of integrated cells. Selection media was replaced daily for five total days of selection. After selection, cells were returned to DMEM + 10% FBS for outgrowth on six-well plates.

After the outgrown cells reached approximately 80% confluence, cells were trypsinized (0.25% Trypsin-EDTA diluted in PBS at a 3 PBS : 2 trypsin ratio) and resuspended in DMEM + 10% FBS supplemented with 100 U/mL penicillin-streptomycin.

Separately, “half and half” conditioned media was made by removing media from a confluent flask of HEK293T cells, filtering it through a 0.22 µm filter, and mixing it 1:1 with fresh media. The resulting conditioned media was supplemented with 100 U/mL penicillin-streptomycin.

Cells were sorted on a Sony MA-900 flow sorter using the gates shown in Figure S7. Briefly, live single cells were identified using forward scatter and side scatter gates, and the mRuby2-positive cells were gated using a roughly rectangular mRuby2-FSC gate. Cells were recovered onto gelatin-coated, pre-warmed plates containing the conditioned media. After a media change and outgrowth in media not containing penicillin-streptomycin, cells were confirmed myco-negative (Lonza MycoAlert).

### Lentiviral integration

#### Lentivirus production

Lenti-X HEK293T cells (Takara Bio, 632180) grown in DMEM + 10% FBS were seeded at 10^6^ cells per well of a 6-well plate. The following day (day 1), 1 µg of the third-generation lentiviral expression plasmid, 1 µg of the packaging plasmid (psPAX2, Addgene #12260), and 2 µg of the envelope plasmid (pMD2.G / VSVG, Addgene #12259) per well were co-transfected using PEI as described above. After 6 hours, the media was aspirated and replaced with 1.25 mL of DMEM + 10% FBS + 25 mM HEPES (Sigma-Aldrich, H3375). On the following day (day 2), the media was collected, stored at 4°C, and replaced with HEPES-buffered DMEM + 10% FBS. On day 3, the media was again collected. The collected media was filtered through a 0.45 µm PES filter.

To the filtered virus-containing media, Lenti-X Concentrator (Takara Bio, 631232) was added in a 3 parts media : 1 part concentrator volume ratio, mixed gently, and left overnight at 4°C. On day 4, the media was centrifuged at 1500 rcf at 4°C for 45 minutes. The supernatant was aspirated, and the resulting pellet was resuspended to a total volume of 200 µL in HEPES-buffered DMEM + 10% FBS. Virus was used immediately or stored at −80°C.

#### Lentivirus titration

Regularly passaged HEK293T cells were seeded at a concentration of 15,000 cells per well of a 96-well plate in DMEM + 10% FBS on the day of the transduction. Cells were combined with 5 µg/mL polybrene (hexadimethrine bromide, Sigma-Aldrich, H9268-5G) and a two-fold serial dilution of the produced virus (highest concentration: 5.0 µL concentrated virus per well). The resulting cell, polybrene, and virus mixture was plated onto 96-well plates coated with 0.1% gelatin. Three days later, the resulting cells were dissociated using 0.25% Trypsin-EDTA diluted in PBS at a 3 PBS : 2 trypsin ratio, and data was collected using an Attune NxT flow cytometer. Single, live cells were selected using forward scatter and side scatter gates. Transduced cells were identified using an mRuby2 gate that excluded untransduced cells.

#### HEK293T transduction

Regularly passaged HEK293T cells were seeded on the day of viral transduction in suspension at 150,000 cells per 12-well. Each 12-well was transduced with concentrated lentivirus produced from 6-well plate and titered to have a multiplicity of infection (MOI) of 2. Fresh DMEM + 10% FBS was included to reach a final volume of 2 mL per 12-well, and 5 µg/mL polybrene was added to increase transduction efficiency. Three days later, the resulting cells were dissociated and labeled for HCR Flow-FISH.

### HCR RNA-FISH

In all HCR RNA-FISH experiments here, we use Molecular Instruments probe sets for mRuby and tagBFP compatible with B2 amplifiers conjugated to Alexa Fluor™ 514. The FISH protocol as well as hybridization and wash buffer compositions were based on those reported by Choi *et al.* and modified to improve cell recovery for flow cytometry [27]. Amplification buffer composition was based on that reported by Jia *et al* [60]. All buffer compositions are reported in Table S5.

#### FISH imaging

Cells grown on Geltrex-coated glass-bottom plates were fixed by incubating with 4% paraformaldehyde (PFA) solution (EMD Millipore, 818715) for one hour at 4°C. After washing the cells three times with cold PBS, the cells were permeabilized using 0.5% Tween-20 (Sigma-Aldrich, P2287) overnight at 4°C. Next, cells were washed twice with 2X SSC and then incubated with hybridization buffer for 30 minutes at 37°C. Probe set stock solution was diluted to 4 nM in hybridization buffer. Cells were incubated in this probe solution overnight at 37°C.

Following hybridization, cells were incubated with wash buffer for five minutes at 37°C, and this was repeated for a total of four washes. Then, cells were incubated with 5X SSCT for five minutes at room temperature, and this was repeated for a total of two washes. After these washes, cells were incubated in amplification buffer for 30 minutes at room temperature. Amplifier solution was prepared by combining separately snap-cooled hairpins h1 and h2 at a concentration of 60 nM in amplification buffer. Cells were incubated with this amplifier solution overnight at room temperature.

Following amplification, cells were incubated with 5X SSCT for five minutes at room temperature, and this was repeated for a total of five washes. Finally, wells were filled with PBS for imaging using a Nikon Ti2-E fluorescence microscope.

#### HCR Flow-FISH

After suspension in PBS, cells were transferred to 96-well v-bottom plate for HCR Flow-FISH. After each resuspension, spins were performed at 500 rcf for 5 minutes with default settings, unless otherwise noted. Cells were first fixed through incubation in 4% PFA for 15 minutes at room temperature. After spinning, cells were then permeabilized using 0.5% Tween-20 for 15 minutes at room temperature. Next, cells were spun and resuspended in hybridization buffer for 30 minutes at 37°C. During this incubation, probe set stock solution was diluted in hybridization buffer to a concentration of 14 nM for transfected cells or 28 nM for integrated cell lines. Cells were spun and resuspended in this probe solution for incubation overnight at 37°C. Due to the viscosity of the hybridization buffer, these spins were performed with reduced deceleration speed to minimize cell loss.

Following hybridization, cells were spun and resuspended in wash buffer for 15 minutes at 37°C. Then, cells were spun and resuspended in 5X SSCT for five minutes at room temperature. After these washes, cells were spun and resuspended in amplification buffer for 30 minutes at room temperature. Amplifier solution was prepared by combining separately snap-cooled hairpins h1 and h2 at a concentration of 130 nM in amplification buffer. Cells were spun and then incubated with this amplifier solution overnight at room temperature.

Following amplification, cells were spun and resuspended in 5X SSCT for one 30 minute incubation and one five minute incubation at room temperature. Finally, cells were spun and resuspended in PBS for flow cytometry.

### Flow cytometry

All flow cytometry data was collected using an Attune NxT flow cytometer with channel mappings and voltages reported in Table S6. Data from HCR-Flow-FISH experiments was compensated using the matrix reported in Table S7. Single cells were selected using forward scatter and side scatter gates. Transfected cells were gated based on expression of a co-transfected marker. For lentiviral transduction, the top 85% of cells were gated based on mRuby2 expression, assuming a Poisson distribution of integration events corresponding to an MOI of 2.

### RT-qPCR

For concurrent RT-qPCR and HCR Flow-FISH analysis, PiggyBac-integrated cell lines were plated in biological triplicate at a density of 300,000 cells per well in 6-well plates. After three days of growth, cells were dissociated by adding 0.25% Trypsin-EDTA (diluted in PBS) for four minutes, followed by quenching with an equal volume of DMEM + 10% FBS. Each sample of dissociated cells was split in half, with equal amounts going to HCR Flow-FISH processing or RNA isolation. RNA was isolated using the Monarch Total RNA Miniprep Kit (New England Biolabs, T2010) with an additional Dnase I (New England Biolabs, M0570) treatment step. RNA samples were eluted into 50 µL of nuclease-free water. cDNA was synthesized from 6 µL of eluted RNA using the ProtoScript First Strand cDNA Synthesis Kit (New England Biolabs, E6300) with oligo-dT primers. cDNA samples were stored at −20°C until qPCR.

qPCR was performed at the MIT BioMicro Center on a Roche LightCycler 480 with four technical replicates per condition. Reaction mixes were assembled using 2.5 µL KAPA SYBR FAST qPCR Master Mix (2X) Universal (Kapa Biosystems, KK4600), 0.5 µL 2 µM forward and reverse primers, 0.5 µL cDNA product, and 1.5 µL nuclease-free water. The primer sequences used for each gene are reported in Table S8. Using the “High Sensitivity” analysis mode, *C_t_* values were called. Pooling over technical replicates by taking the mean, Δ*C_t_* values were calculated relative to the GADPH levels for each sample and used to calculate relative expression levels.

### Long-read sequencing

RNA from polyclonal PiggyBac-integrated cell lines was isolated using the Monarch Total RNA Miniprep Kit with an additional Dnase I treatment step. RNA samples were eluted into 50 µL of nuclease-free water.

Direct RNA sequencing was performed on an Oxford Nanopore GridION device using the Direct Sequencing Kit (SQK-RNA004, date accessed 15 May 2024), MinION RNA flow cell (FLO-MIN00RA), and data pre-processing was performed with MinKNOW (v24.06.10). Libraries were constructed individually with the following modifications to optimize fragment yield and quantity: (1) approximately 1.2 µg of total RNA was used in 8 µL total volumes; and (2) All binding and elution steps were doubled, with a minimum bead binding time of 5 minutes. Basecalling was performed on-device using the ‘super-accurate basecalling’ model in dorado version 7.4.12. The resulting .fastq files were aligned using minimap2 version 2.26 (flags: -ax splice -uf -k14) to custom human reference genomes combining GRCh38 v108 with the plasmid sequence for each construct.

Unique reads mapping to the construct sequences were isolated using bedtools. Major isoform start sites were manually identified from the reads looking at a density distribution of read starts. For any analyses quantifying the start or end positions of the reads, reads were filtered to remove any reads whose 5’ end was greater than 25 nt away from major isoform start sites to avoid any artifacts introduced by 5’ truncation prevalent in long-read RNA sequencing data. Major transcription start sites (TSS) and intron locations for each promoter sequence are reported in Table S1.

Differential expression of endogenous genes was analyzed by comparing data from each cell line to data from the cell line integrated with hPGK-mRuby2-P2A-PuroR-bGH, which has the weakest transgene expression. This analysis assumes the hPGK promoter has the fewest off-target effects on endogenous gene expression. For each gene, the fold change in the read counts per million (CPM) relative to the hPGK baseline was calculated, and any fold change with an absolute value greater than 1.5 was considered different from the baseline.

