## Supplementary Figures for "High-resolution profiling reveals coupled transcriptional and translational regulation of transgenes"

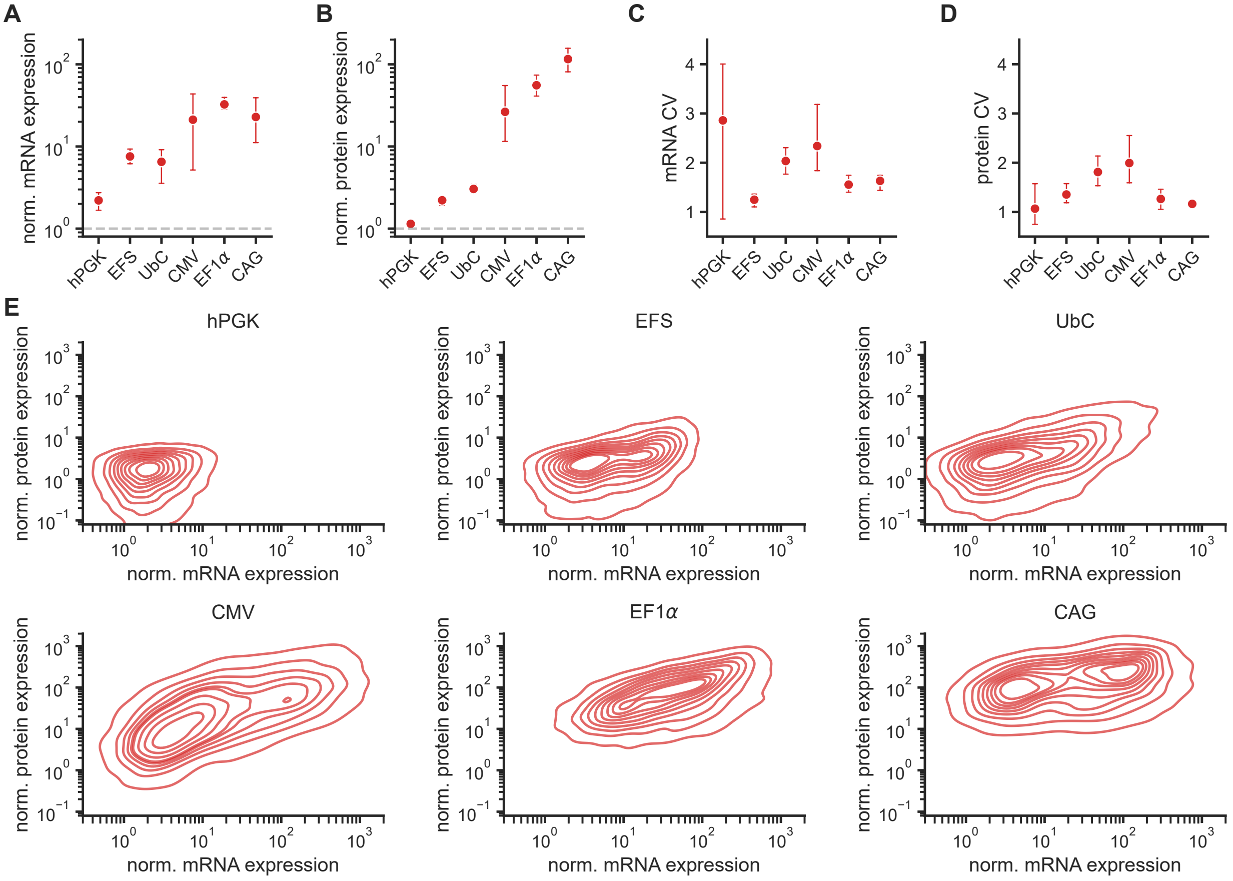


**Figure S1. Additional mRuby2 transfection characterization results.**
**A**. Normalized mRuby2 mRNA expression in HEK293T transfection with varying promoter sequences. Dashed line indicates background fluorescence level.
**B**. Normalized mRuby2 protein expression in HEK293T transfection with varying promoter sequences. Dashed line indicates background fluorescence level.
**C**. mRuby2 mRNA coefficient of variation in HEK293T transfection with varying promoter sequences.
**D**. mRuby2 protein coefficient of variation in HEK293T transfection with varying promoter sequences.
**E**. Joint distributions of mRuby2 mRNA and protein fluorescence in HEK293T transfection with varying promoter sequences. Data was randomly downsampled to 10,000 cells per condition for plotting. Normalized expression is calculated as the fold change of fluorescence intensity relative to a non-transfected sample. Points represent means of three biological replicates, and error bars represent the 95% confidence interval. All data is in arbitrary units from a flow cytometer.

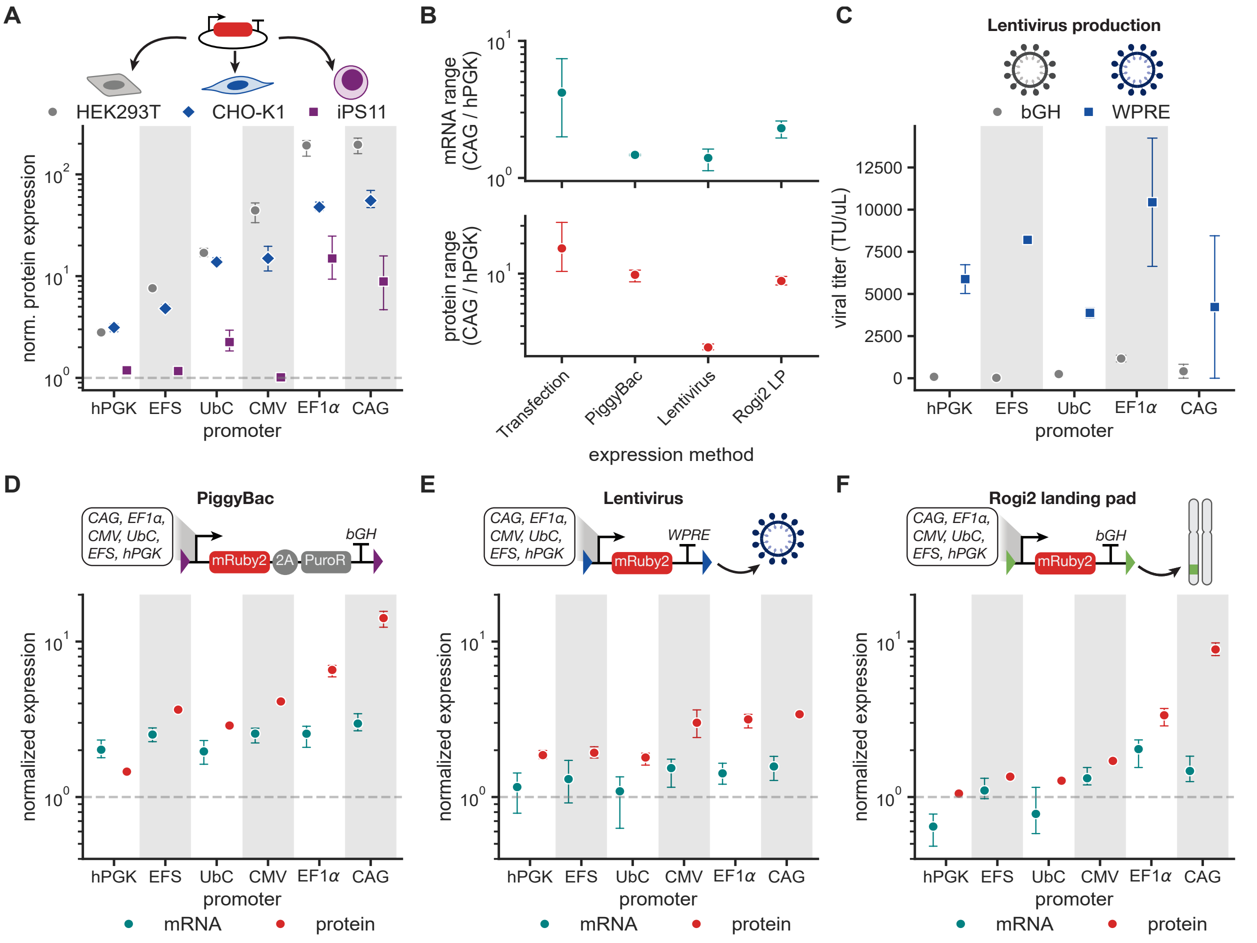


**Figure S2. Generalizability of promoter strength across cell types and integration contexts.**
**A**. Normalized protein expression for HEK293T, CHO-K1, and iPS11 cells transfected with a plasmid encoding mRuby2 driven by six different constitutive promoters with a bGH poly(A) tail.
**B**. Range of mRNA and protein expression in different expression contexts as defined by the ratio between expression with the strong promoter CAG and the weak promoter hPGK.
**C**. Mean viral titer in transducing units (TU) per µL of lentivirus produced with varying promoter and 3' UTR sequence. Points represent the mean ± the 95% confidence interval for two batches of virus.
**D**. Normalized protein and mRNA expression for HEK293T cells PiggyBac-integrated with mRuby2-2A-PuroR-bGH driven by six different constitutive promoters.
**E**. Normalized protein and mRNA expression for HEK293T cells Lentivirus-integrated with mRuby2-WPRE driven by six different constitutive promoters.
**F**. Normalized protein and mRNA expression for Rogi2 LP HEK293T cells site-specifically integrated with mRuby2-bGH driven by six different constitutive promoters.
Normalized expression is calculated as the fold change of fluorescence intensity relative to a non-transfected sample. Dashed lines represent background fluorescence level of a non-transfected sample. Points represent the mean of three biological replicates ± the 95% confidence interval. All data is in arbitrary units from a flow cytometer.

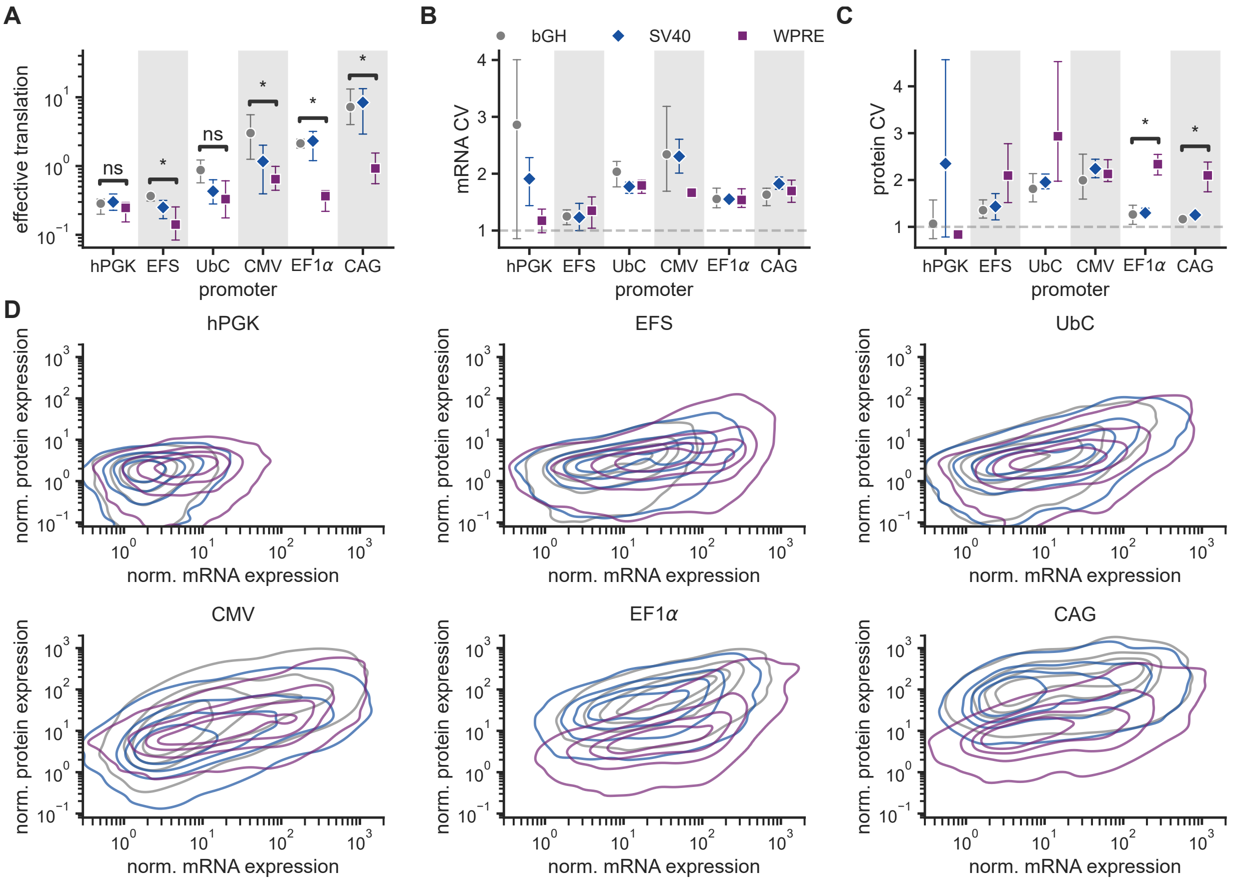


**Figure S3. Additional 3' UTR transfection characterization results.**
**A**. Effective translation rate as calculated by the slope of a line fitted the binned data in Figure 3D. *: $p \leq 0.05$, one-sided Mann-Whitney test
**B**. mRuby2 mRNA coefficient of variation in HEK293T transfection with varying promoter and PAS or 3' UTR sequences.
**C**. mRuby2 protein coefficient of variation in HEK293T transfection with varying promoter and PAS or 3' UTR sequences. *: $p \leq 0.05$, one-sided Mann-Whitney test
**D**. Joint distributions of mRuby2 mRNA and protein fluorescence in HEK293T transfection with varying promoter and PAS or 3' UTR sequences. Data was randomly downsampled to 10,000 cells per condition for plotting.
Normalized expression is calculated as the fold change of fluorescence intensity relative to a non-transfected sample. Points represent means of three biological replicates, and error bars represent the 95% confidence interval. All data is in arbitrary units from a flow cytometer.

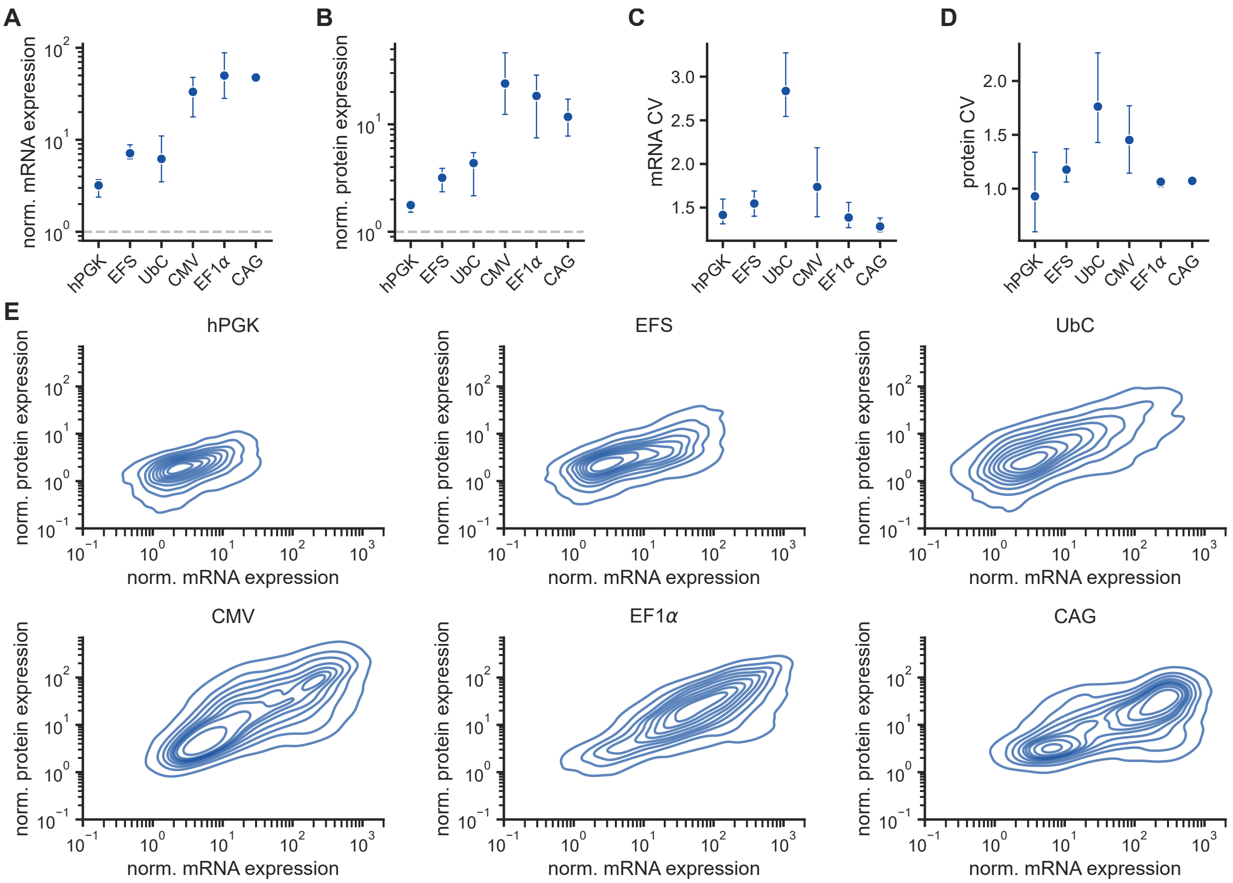


**Figure S4. Additional tagBFP transfection characterization results.**
**A**. Normalized tagBFP mRNA expression in HEK293T transfection with varying promoter sequences. Dashed line indicates background fluorescence level.
**B**. Normalized tagBFP protein expression in HEK293T transfection with varying promoter sequences. Dashed line indicates background fluorescence level.
**C**. TagBFP mRNA coefficient of variation in HEK293T transfection with varying promoter sequences.
**D**. TagBFP protein coefficient of variation in HEK293T transfection with varying promoter sequences.
**E**. Joint distributions of tagBFP mRNA and protein fluorescence in HEK293T transfection with varying promoter sequences. Data was randomly downsampled to 10,000 cells per condition for plotting.
Normalized expression is calculated as the fold change of fluorescence intensity relative to a non-transfected sample. Points represent means of three biological replicates, and error bars represent the 95% confidence interval. All data is in arbitrary units from a flow cytometer.

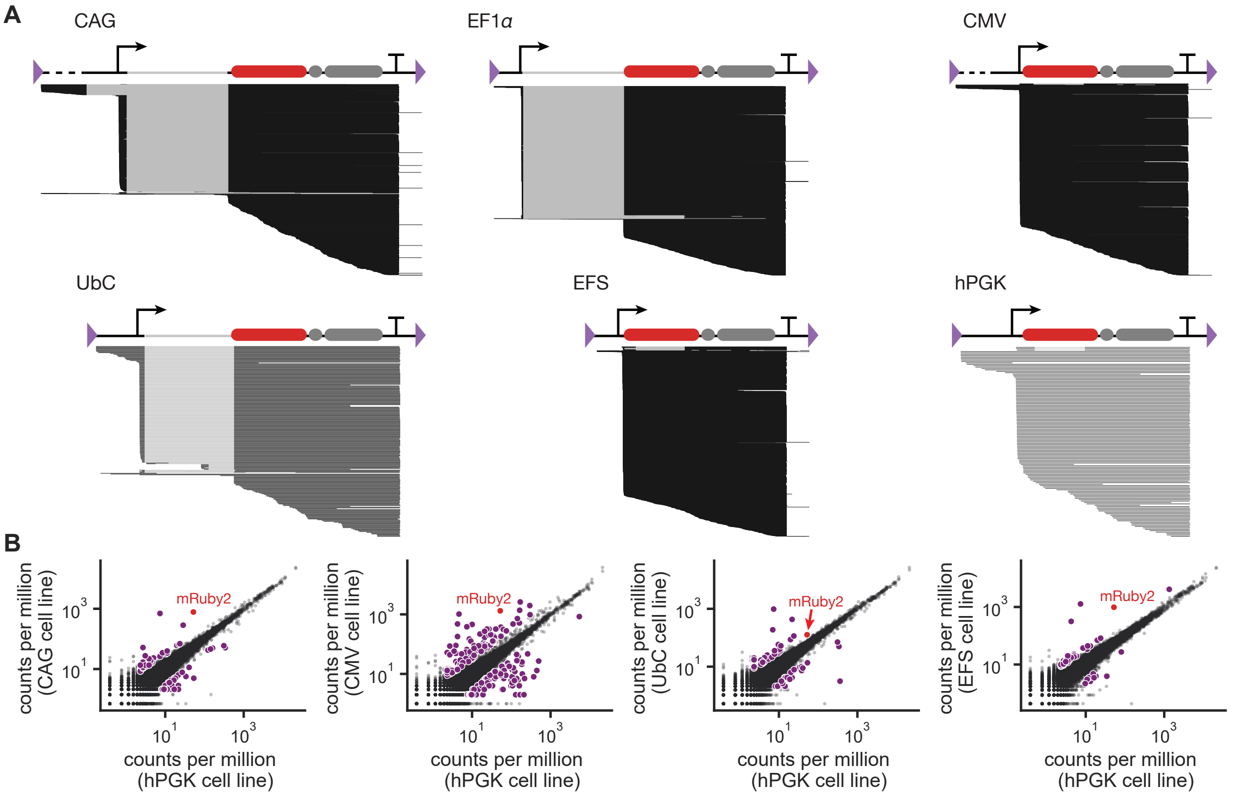


**Figure S5. Additional long-read sequencing results.**
**A**. RNA transcript maps for the six constitutive promoters tested without downsampling.
**B**. Comparison of endogenous gene expression levels between cell lines integrated with the CAG, CMV, UbC, or EFS promoters relative to the cell line integrated with the hPGK promoter. Genes are considered "differentially expressed" between the cell lines if the absolute value of the fold change is greater than 1.5. These genes are indicated in purple. mRuby2 expression is indicated in red.

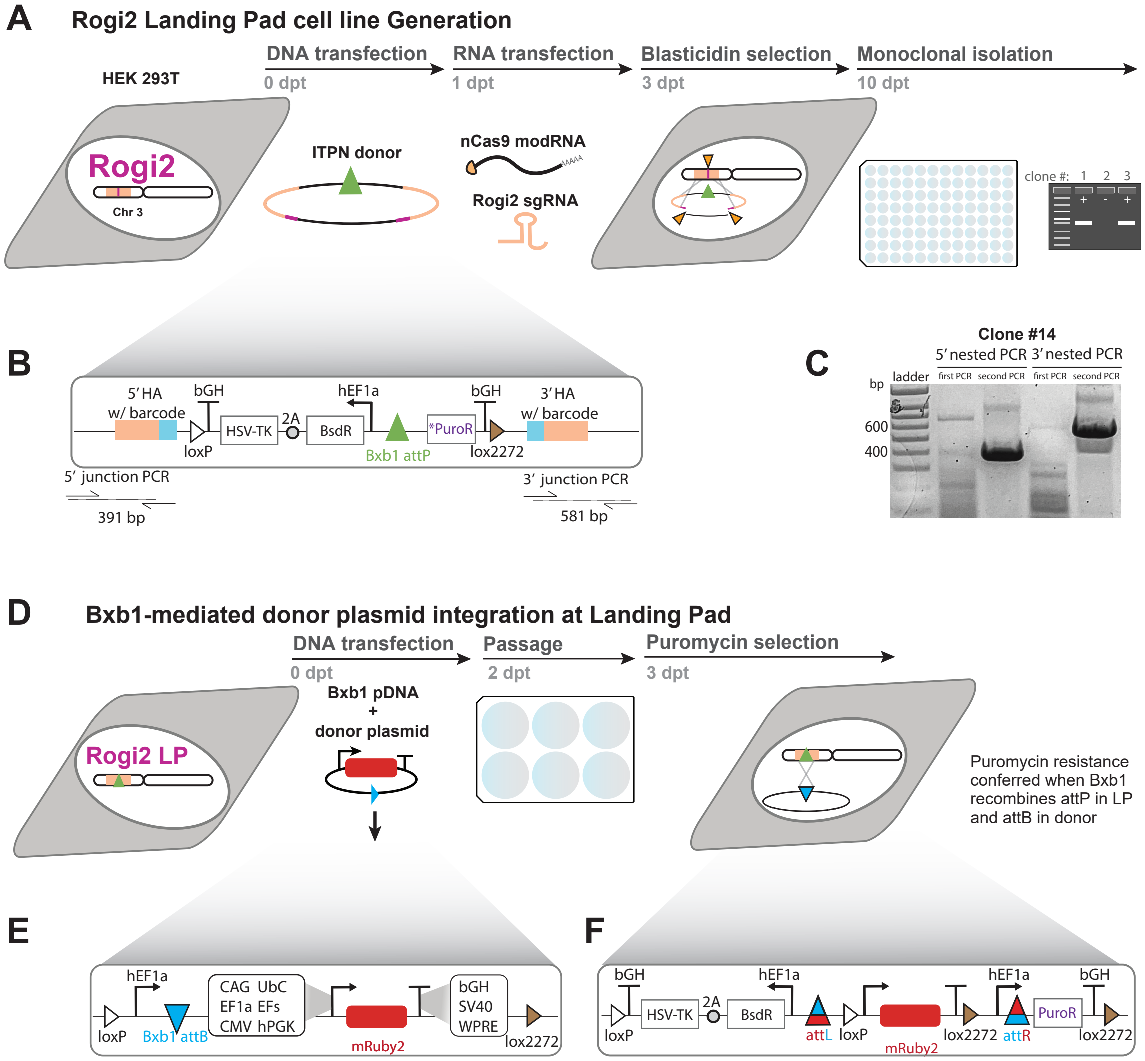


**Figure S6. HEK293T Rogi2 landing pad for evaluating genetic parts in site-specific integration.**
**A**. Workflow for installation of landing pad at Rogi2 via In Trans Paired Nicking (ITPN).
**B**. Architecture of landing pad encoded on Rogi2-specific ITPN donor. The barcodes directly internal to the homology arms are used in 5' and 3' junction PCRs when genotyping candidate clones.
**C**. Nested PCR results for 5' and 3' junctions in monoclone #14, verifying installation of donor DNA at Rogi2.
**D**. Workflow for integrating donor plasmids at Rogi2 landing pad via Bxb1.
**E**. Depiction of genetic components encoded on Bxb1 donor plasmids to integrate the complete set of constitutive promoter-polyA mRuby2 transcriptional units.
**F**. Depiction of genomic sequence after Bxb1-mediated insertion of the donor plasmid at the Landing Pad. The insertion of the hEF1a promoter and start codon in-frame with the Puromycin gene pre-installed at the landing pad confers resistance only to successfully-recombined cells.

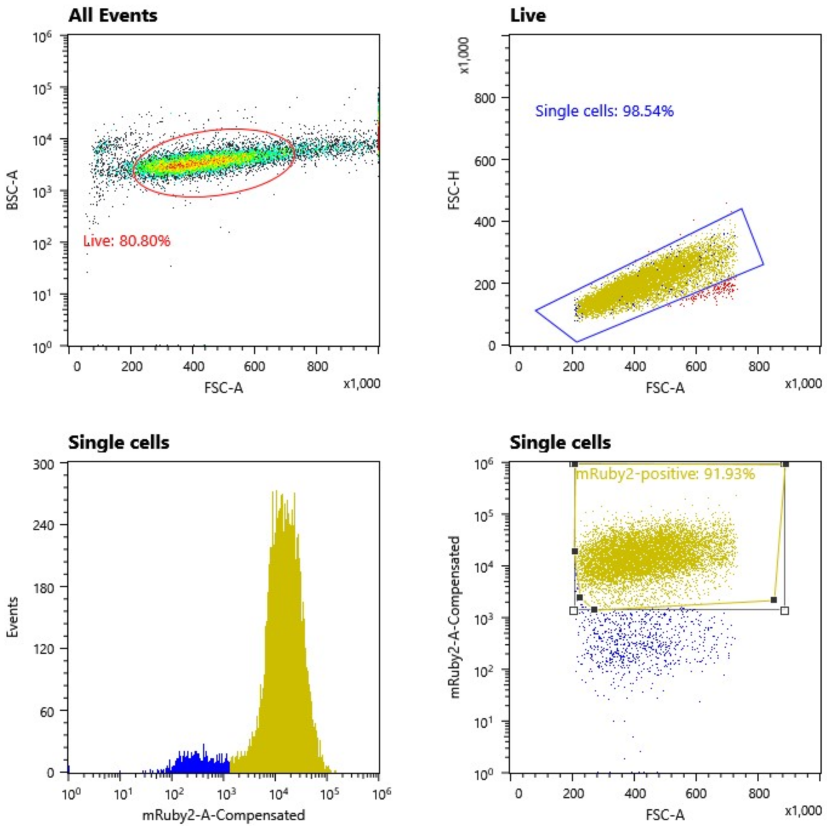


**Figure S7. Gating strategy used for sorting PiggyBac-integrated cell lines on a Sony MA-900 flow sorter.**

### Supplementary Tables

**Table S1.** Constitutive promoter sequences annotated with the most common TSS (bold and underline) and intron (italic and lowercase) coordinates as determined by long-read sequencing

| Promoter | Sequence |
|---|---|
| CAG | TTATTAATAGTAATCAATTACGGGGTCATTAGTTCATAGCCCATATATGGAGTTCCGCGT<br>TACATAACTTACGGTAAATGGCCCGCCTGGCTGACCGCCCAACGACCCCCGCCCATTGAC<br>GTCAATAATGACGTATGTTCCCATAGTAACGCCAATAGGGACTTTCCATTGACGTCAATG<br>GGTGGAGTATTTACGGTAAACTGCCCACTTGGCAGTACATCAAGTGTATCATATGCCAAG<br>TACGCCCCCTATTGACGTCAATGACGGTAAATGGCCCGCCTGGCATTATGCCCAGTACAT<br>GACCTTATGGGACTTTCCTACTTGGCAGTACATCTACGTATTAGTCATCGCTATTACCAT<br>GGTCGAGGTGAGCCCCACGTTCTGCTTCACTCTCCCCATCTCCCCCCCCTCCCCACCCCC<br>AATTTTGTATTTATTTATTTTTTAATTATTTTGTGCAGCGATGGGGGCGGGGGGGGGGGG<br>GGGGCGCGCGCCAGGCGGGGCGGGGCGGGGCGAGGGGCGGGGCGGGGCGAGGCGGAGAGG<br>TGCGGCGGCAGCCAATCAGAGCGGCGCGCTCCGAAAGTTTCCTTTTATGGCGAGGCGGCG<br>GCGGCGGCGGCCCTATAAAAAGCGAAGCGCGCGGCGGGCGGGAGTCGCTGCGCGCTG**C**CT<br>TCGCCCCGTGCCCCGCTCCGCCGCCGCCTCGCGCCGCCCGCCCCGGCTCTGACTGACCGC<br>GTTACTCCCACAG*gtgagcgggcgggacggcccttctcctccgggctgtaattagcgctt*<br>*ggtttaatgacggcttgtttcttttctgtggctgcgtgaaagccttgaggggctccggga*<br>*gggccctttgtgcggggggagcggctcggggggtgcgtgcgtgtgtgtgtgcgtggggag*<br>*cgccgcgtgcggctccgcgctgcccggcggctgtgagcgctgcgggcgcggcgcggggct*<br>*ttgtgcgctccgcagtgtgcgcgaggggagcgcggccgggggcggtgccccgcggtgcgg*<br>*ggggggctgcgaggggaacaaaggctgcgtgcggggtgtgtgcgtgggggggtgagcagg*<br>*gggtgtgggcgcgtcggtcgggctgcaaccccccctgcacccccctccccgagttgctga*<br>*gcacggcccggcttcgggtgcggggctccgtacggggcgtggcgcggggctcgccgtgcc*<br>*gggcggggggtggcggcaggtgggggtgccgggcggggcggggccgcctcgggccgggga*<br>*gggctcgggggaggggcgcggcggcccccggagcgccggcggctgtcgaggcgcggcgag*<br>*ccgcagccattgccttttatggtaatcgtgcgagagggcgcagggacttcctttgtccca*<br>*aatctgtgcggagccgaaatctgggaggcgccgccgcaccccctctagcgggcgcggggc*<br>*gaagcggtgcggcgccggcaggaaggaaatgggcggggagggccttcgtgcgtcgccgcg*<br>*ccgccgtccccttctccctctccagcctcggggctgtccgcgggggggacggctgccttcg*<br>*ggggggacggggcagggcggggttcggcttctggcgtgtgaccggcggctctagagcctc*<br>*tgctaaccatgttcatgccttcttctttttcctacag*CTCCTGGGCAACGTGCTGGTTAT<br>TGTGCTGTCTCATCATTTTGGCAAA |

Continued on next page

**Table S1 – continued from previous page**

| Promoter | Sequence |
|---|---|
| EF1α | GCTCCGGTGCCCGTCAGTGGGCAGAGCGCACATCGCCCACAGTCCCCGAGAAGTTGGGGG<br>GAGGGGTCGGCAATTGAACCGGTGCCTAGAGAAGGTGGCGCGGGGTAAACTGGGAAAGTG<br>ATGTCGTGTACTGGCTCCGCCTTTTTCCCGAGGGTGGGGGAGAACCGTATATAAGTGCAG<br>TAGTCGCCGTGAACGTTCTTTTTCG**C**AACGGGTTTGCCGCCAGAACACAG*gtaagtgccg*<br>*tgtgtggttcccgcgggcctggcctctttacgggttatggcccttgcgtgccttgaatta*<br>*cttccacgcccctggctgcagtacgtgattcttgatcccgagcttcgggttggaagtggg*<br>*tgggagagttcgaggccttgcgcttaaggagccccttcgcctcgtgcttgagttgaggcc*<br>*tggcttgggcgctggggccgccgcgtgcgaatctggtggcaccttcgcgcctgtctcgct*<br>*gctttcgataagtctctagccatttaaaatttttgatgacctgctgcgacgctttttttc*<br>*tggcaagatagtcttgtaaatgcgggccaagatctgcacactggtatttcggtttttggg*<br>*gccgcgggcggcgacggggcccgtgcgtcccagcgcacatgttcggcgaggcggggcctg*<br>*cgagcgcggccaccgagaatcggacgggggtagtctcaagctggccggcctgctctggtg*<br>*cctggcctcgcgccgccgtgtatcgccccgccctgggcggcaaggctggcccggtcggca*<br>*ccagttgcgtgagcggaaagatggccgcttcccggccctgctgcagggagctcaaaatgg*<br>*aggacgcggcgctcgggagagcgggcgggtgagtcacccacacaaaggaaaagggccttt*<br>*ccgtcctcagccgtcgcttcatgtgactccacggagtaccgggcgccgtccaggcacctc*<br>*gattagttctcgagcttttggagtacgtcgtctttaggttggggggaggggttttatgcg*<br>*atggagtttccccacactgagtgggtggagactgaagttaggccagcttggcacttgatg*<br>*taattctccttggaatttgccctttttgagtttggatcttggttcattctcaagcctcag*<br>*acagtggttcaaagtttttttcttccatttcag*GTGTCGTGAG |
| CMV | CGTTACATAACTTACGGTAAATGGCCCGCCTGGCTGACCGCCCAACGACCCCCGCCCATT<br>GACGTCAATAATGACGTATGTTCCCATAGTAACGCCAATAGGGACTTTCCATTGACGTCA<br>ATGGGTGGAGTATTTACGGTAAACTGCCCACTTGGCAGTACATCAAGTGTATCATATGCC<br>AAGTACGCCCCCTATTGACGTCAATGACGGTAAATGGCCCGCCTGGCATTATGCCCAGTA<br>CATGACCTTATGGGACTTTCCTACTTGGCAGTACATCTACGTATTAGTCATCGCTATTAC<br>CATGGTGATGCGGTTTTGGCAGTACATCAATGGGCGTGGATAGCGGTTTGACTCACGGGG<br>ATTTCCAAGTCTCCACCCCATTGACGTCAATGGGAGTTTGTTTTGGCACCAAAATCAACG<br>GGACTTTCCAAAATGTCGTAACAACTCCGCCCCATTGACGCAAATGGGCGGTAGGCGTGT<br>ACGGTGGGAGGTCTATATAAGCAGAGCTGAGCTCGTTTAGTGAACCGTC**A**GATCGCCTGG<br>AGACGCCATCCACGCTGT |

Continued on next page

**Table S1 – continued from previous page**

| Promoter | Sequence |
|---|---|
| UbC | GGCCTCCGCGCCGGGTTTTGGCGCCTCCCGCGGGCGCCCCCCTCCTCACGGCGAGCGCTG<br>CCACGTCAGACGAAGGGCGCAGCGAGCGTCCTGATCCTTCCGCCCGGACGCTCAGGACAG<br>CGGCCCGCTGCTCATAAGACTCGGCCTTAGAACCCCAGTATCAGCAGAAGGACATTTTAG<br>GACGGGACTTGGGTGACTCTAGGGCACTGGTTTTCTTTCCAGAGAGCGGAACAGGCGAGG<br>AAAAGTAGTCCCTTCTCGGCGATTCTGCGGAGGGATCTCCGTGGGGCGGTGAACGCCGAT<br>GATTATATAAGGACGCGCCGGGTGTGGCACAGCTAGTTCCGT**C**GCAGCCGGGATTTGGGT<br>CGCGGTTCTTGTTTGTGGATCGCTGTGATCGTCACTTG*gtgagtagcgggctgctgggct*<br>*ggccggggctttcgtggccgccgggccgctcggtgggacggaagcgtgtggagagatcgc*<br>*caagggctgtagtctgggtccgcgagcaaggttgccctgaactgggggttggggggagcg*<br>*cagcaaaatggcggctgttcccgagtcttgaatggaagacgcttgtgaggcgggctgtga*<br>*ggtcgttgaaacaaggtgggggcatggtgggcggcaagaacccaaggtcttgaggcctt*<br>*cgctaatgcgggaaagctcttattcgggtgagatgggctggggcaccatctggggaccct*<br>*gacgtgaagtttgtcactgactggagaactcggtttgtcgtctgttgcgggggcggcagt*<br>*tatggcggtgccgttgggcagtgcacccgtacctttgggagcgcgcgccctcgtcgtgtc*<br>*gtgacgtcacccgttctgttggcttataatgcagggtggggccacctgtcggtaggtgtg*<br>*cggtaggcttttctccgtcgcaggacgcagggttcgggcctagggtaggctctcctgaat*<br>*cgacaggcgccggacctctggtgaggggagggataagtgaggcgtcagtttctttggtcg*<br>*gttttatgtacctatcttcttaagtagctgaagctccggttttgaactatgcgctcgggg*<br>*ttggcgagtgtgttttgtgaagtttttaggcaccttttgaaatgtaatcatttgggtca*<br>*atatgtaattttcagtgttagactagtaaattgtccgctaaattctggccgtttttggct*<br>*tttttgttag*AC |
| EFS | GGGCAGAGCGCACATCGCCCACAGTCCCCGAGAAGTTGGGGGGAGGGGTCGGCAATTGAA<br>CCGGTGCCTAGAGAAGGTGGCGCGGGGTAAACTGGGAAAGTGATGTCGTGTACTGGCTCC<br>GCCTTTTTCCCGAGGGTGGGGGAGAACCGTATATAAGTGCAGTAGTCGCCGTGAACGTTC<br>TTTTTCG**C**AACGGGTTTGCCGCCAGAACACAGT |
| hPGK | GGGGTTGGGGTTGCGCCTTTTCCAAGGCAGCCCTGGGTTTGCGCAGGGACGCGGCTGCTC<br>TGGGCGTGGTTCCGGGAAACGCAGCGGCGCCGACCCTGGATCTCGCACATTCTTCACGTC<br>CGTTCGCAGCGTCACCCGGATCTTCGCCGCTACCCTTGTGGGCCCCCCGGCGACGCTTCC<br>TGCTCCGCCCCTAAGTCGGGAAGGTTCCTTGCGGTTCGCGGCGTGCCGGACGTGACAAAC<br>GGAAGCCGCACGTCTCACTAGTACCCTCGCAGACGGACAGCGCCAGGGAGCAATGGCAGC<br>GCGCCGACCGCGATGGGCTGTGGCCAATAGCGGCTGCTCAGCAGGGCGCGCCGAGAGCAG<br>CGGCCGGGAAGGGGCGGTGCGGGAGGCGGGGTGTGGGGCGGTAGTGTGGGCCCTGTTCCT<br>GCCCGCGCGGTGTTCCGCATTCTGCAAGC**C**TCCGGAGCGCACGTCGGCAGTCGGCTCCCT<br>CGTTGACCGAATCACCGACCTCTCTCCCCAGA |

**Table S2.** Differentially expressed genes reported in Figure 5J for PiggyBac-integrated cell lines with the indicated promoters.

| Integrated promoter | Differentially expressed genes |
|---|---|
| CAG | R3HDM1, PIK3CB, CAMK2B, NGEF, ATRX, KIAA0391, MID1, EYA1, ACBD5, ACAD10, BCHE, FGF12, TROVE2, KIF17, VAMP8, DOCK6, SERPINF1, TESMIN, BEX1, LMO7, ALG10, SCAPER, TBCD, NR6A1, ZKSCAN2, SIM2, ANTXR2, LARGE2, PRELID1, STAT2, SNHG11, RALGAPA1, SLC29A2, FBXW8, HASPIN, BX322650.1, NELL2, ZBTB40, ATL1, ZNF536, FAM169A, SOWAHA, RNY1, RNA5SP202, SNORA2B, MT-TW, MT-TS1, CSKMT, NMD3P1, MT-ATP8, MTCO2P12, LINC-PINT, RPL3P4, SHANK3, AP001994.1, SNHG25, AC010328.2, FENDRR, AC010422.3, AC009275.1, CU633904.1, RNA5-8SN1, RNA5-8SN2, FP236383.2, SCARNA4, AL772307.1 |
| EF1α | AASS, YIPF1, DGKA, MTMR2, RBFOX2, MID1, MAN2B1, ADAP1, APBB3, CEP104, KIF17, FBXO30, G0S2, TMEM255A, TBC1D5, BEX1, DTNA, PTGR2, INTS4, INCENP, EXO5, KATNAL2, KIF5C, KCTD19, ZNF692, RALGAPA1, WDR25, HASPIN, MEIOC, INCA1, PPP1R26, CACNA1H, TCF4, SPG7, FAM169A, SNORA33, MT-TW, MT-TS1, AL031133.1, NMD3P1, MT-ATP8, MTCO2P12, BDNF-AS, GMDS-DT, SCARNA21, TMPO-AS1, AC015917.2, AC015813.2, AC010328.2, AC009275.1, RNA5-8SN2, FP236383.2, SCARNA4 |
| UbC | ALS2, TYMP, CAMK2B, GRIPAP1, RFFL, KIAA0391, MID1, TAF6, KANK1, LIFR, TROVE2, DCAF4, ST3GAL3, TBC1D5, INPP5K, DTNA, ITGA7, ZFHX3, AZIN2, ANK2, SLC16A2, INTS4, CAMK4, VPS39, CEP120, ERCC6L2, OSBP2, ZNF286A, ZNF790, SPG7, TMEM116, FAM169A, RNY1, PRR13, MT-TS1, MT-TK, LAMTOR5-AS1, SATB1-AS1, UBE2SP1, MTCO1P12, PTPRG-AS1, GMDS-DT, AP002748.3, AC008915.2, AL118516.1, AC009275.1, INO80B-WBP1, RNA5-8SN1, RNA5-8SN2, SCARNA4, AL772307.1 |
| EFS | TYMP, NGEF, SYT1, WIPI1, ANO8, MTMR2, CCNK, KIAA0391, MID1, CCDC113, TBC1D9, KIF17, G0S2, DOCK6, STK33, SH3BP5, SBF2, LMO7, GATA4, TMEM8B, SLC16A2, DGKE, ZKSCAN2, ANTXR2, APBB2, LARGE2, ZBTB39, KCTD19, ZNF692, FAM222B, INSM1, CCDC106, RALGAPA1, FBXW8, FAM210A, MEIOC, CCDC151, TMEM116, ATL1, FAM169A, MT-ND4, SNORA33, TRIM39, MT-TS1, LAMTOR5-AS1, AL133351.1, MTCO1P12, SCARNA18B, SCARNA4 |

Continued on next page

**Table S2 – continued from previous page**

| Integrated promoter | Differentially expressed genes |
|---|---|
| CMV | STPG1, ZNF263, CYTH3, PHF7, RCN1, NEDD4L, YBX3, EIF4B, DHX8, REEP1, TMEM260, PABPC1, IGF2BP2, ENO1, WDR62, PAK3, TULP3, VDAC3, RUNX1T1, SLC1A3, HADHA, CHMP5, MTMR2, L2HGDH, LZTS3, IGBP1, CCNK, SLC22A17, HSP90AB1, SEPT3, ARSA, MID1, CSPP1, STK3, ADAP1, IQCE, SPATA6L, UBE2S, ZPR1, GAPDH, TPI1, TMCO6, RAD50, LIFR, BCHE, MRPL3, SUMO1, TROVE2, VAMP8, ZBTB45, EPC1, TENT4B, PYROXD1, MAPK8IP1, SEPT7, NT5C3A, PFKFB2, C1orf61, ZNF133, MAP2K2, TRAF2, ATP1B2, PUDP, STK33, SH3BP5, IMMT, EIF5A, LARGE1, LDHA, DMTF1, EPHA7, GDF11, SKIL, TMOD1, CDK5RAP2, TMEM8B, VARS2, SSB, OLA1, NDUFA9, PARN, NOB1, RPL13A, TPM3, CALM2, SPOPL, HSPD1, ANK2, KDM6A, SNX30, PARD3, ASAP2, ACSL1, PSD3, RPGR, CCNB2, PSMD4, FDPS, PSMC2, RBBP, WDCP, MAD2L1, STEAP1, RAD21, HNRNPK, LARGE2, MMADHC, FNTA, KCTD19, PRELID1, PCDH7, SETMAR, STAT2, TPT1-AS1, MANEA, CCDC106, C11orf80, RALGAPA1, DOLK, BNIP3, WDR25, HASPIN, AP3S1, RPS27, MROH1, SRP9P1, PIPSL, SGSH, ERCC6L2, KPNA2, EWSR1, RGPD6, SMTN, OSBP2, MORF4L1, FSD2, SAMD11, ZNF383, ZNF559, ZBED6CL, PPP1R26, BNIP3P1, COL4A6, SPG7, CCDC151, SH3BGRL2, MTCO3P12, HMGN2, SELENOT, OSTC, RASGEF1A, RNY1, RNA5S2, VTRNA1-3, PRR13, CCDC85C, SMG1P2, HMGN1, RNPS1, MT-TL1, MT-TW, MT-TD, AL445305.1, AL354714.2, LAP3P2, OLA1P1, SUGT1P2, RPL13AP7, AL034370.1, HNRNPA3P3, AL512633.1, MRPL3P1, AL596087.1, MORF4L1P1, RPL21P28, SNORA11B, CCDC18-AS1, RTCA-AS1, TMEM183B, UBE2D3P1, RPS29P3, MTND1P23, UQCRFS1P1, FTCDNL1, RHEBP2, AC092017.1, ZNF717, SPCS2P4, EEF1DP1, NAMPTP1, AC011005.1, LINC-PINT, TAPBP, ST13P4, RPL3P4, SELENOTP1, RPS28, AC099560.2, AC026462.1, SUMO2P1, AC013470.3, KPNA2P1, RPL13AP5, MTCO1P12, PTENP1, AC117409.1, RPL36A, PSMC1P1, GATA2-AS1, RBBP4P1, AC139887.2, RNPS1P1, GMDS-DT, SCARNA8, SCARNA21B, SCARNA11, AC080023.2, EEF1G, AL109766.1, RBM17P4, AF274858.1, AC092115.1, CCPG1, YBX3P1, RNU4ATAC, MIR3685, PTP4A2P1, AC010422.3, AL035530.2, AL135925.1, RNU11, FP236383.2, FP671120.3, SCARNA4, MIR1244-3, AL772307.1, AL591485.1 |

| BsaI Golden Gate cloning | | | |
|---|---|---|---|
| Plasmid | Sequence type | Upstream overhang | Downstream overhang |
| pPV0 | pShip backbone | CGCT | TACT |
| pPV1 | Promoter sequence | TACT | AATG |
| pPV2 | Coding sequence | AATG | CAAC |
| pPV3 | Polyadenylation sequence or 3' UTR | CAAC | CGCT |
| **PaqCI Golden Gate cloning** | | | |
| Plasmid | Sequence type | Upstream overhang | Downstream overhang |
| pShip | Single transcriptional unit | TACT | CGCT |
| pHarbor | Backbone for genomic integration | CGCT | TACT |

**Table S3.** Type IIS restriction enzyme overhangs used for golden gate cloning

| Plasmid | Sequence of interest | Sequence source | Notes |
|---|---|---|---|
| pKG1117 | pPV0-pShip backbone | Addgene #29652 | |
| pKG2039 | pPV1-CAG | Addgene #140534 | Sequence has an internal PaqCI site |
| pKG2019 | pPV1-EF1a | Addgene #138730 | |
| pKG0618 | pPV1-CMV | Addgene #40651 | |
| pKG1988 | pPV1-UbC | Addgene #25734 | Sequence was domesticated by mutating internal BsaI sites |
| pKG1179 | pPV1-EFS | Addgene #138730 | EF1a sequence excluding intron |
| pKG1367 | pPV1-hPGK | Addgene #41393 | Sequence was domesticated by mutating internal BsaI sites |
| pKG2402 | pPV1-TRE3G (Tet-On) | Addgene #63800 | |
| pKG0743 | pPV1-ZF43x6-C (COMET) | Addgene #138732 | |
| pKG3657 | pPV1-ZF10.BSx8 (synZiFTR) | Gift from the Khalil Lab | |
| pKG0586 | pPV2-mRuby2 | Addgene #90236 | |
| pKG2387 | pPV2-mRuby2-P2A-PuroR | Addgene #90236 and #1764 | |
| pKG1055 | pPV2-tagBFP | Addgene #70224 | Sequence includes silent mutations to remove BsaI sites |
| pKG3659 | pPV2-rtTA-P2A-iRFP720 (Tet-On) | Addgene #105840 and gift from the Khalil Lab | |
| pKG3658 | pPV2-NLS-FKBP-ZF43-P2A-NES-VP64-FRB-P2A-iRFP720 (COMET) | Addgene #138844 and #138852 | |
| pKG2197 | pPV2-ZF10-NS3-p65 (synZiFTR 1.0) | Addgene #195468 | |
| pKG0587 | pPV3-bGH | Addgene #105841 | |
| pKG0588 | pPV3-SV40 | Addgene #25734 | |
| pKG1313 | pPV3-WPRE | Addgene #25734 | |
| pKG0893 | pHarbor-LentiX1 backbone | Addgene #17297 | |
| pKG1334 | pHarbor-PiggyBac backbone | Addgene #63800 | |
| pKG3560 | pHarbor-Rogi2 backbone | Addgene #198040 | |

**Table S4.** List of pPV and pHarbor plasmids used in Golden Gate cloning

| Hybridization buffer (make fresh for every use) | | |
|---|---|---|
| Component | Source | Volume |
| 30% formamide | Fisher Scientific, BP227 | 3 mL |
| 20X SSC | Fisher Scientific, S2713 and BP3271 | 2.5 mL |
| 1M citric acid, pH 6 | Fisher Scientific, BP327 and A142-212 | 90 µL |
| 10% Tween-20 | Sigma-Aldrich, P2287 | 100 µL |
| 50X Denhardt's solution | Fisher Scientific, AAJ63135AD | 200 µL |
| 50% dextran sulfate | Fisher Scientific, BP1585 | 1.5 mL |
| 20 mg/mL BSA | Sigma-Aldrich, A2058 | 50 µL |
| nuclease-free water | | to 10 mL final volume |
| **Wash buffer (store at -20°C for up to one week)** | | |
| Component | Source | Volume |
| 30% formamide | Fisher Scientific, BP227 | 3 mL |
| 20X SSC | Fisher Scientific, S2713 and BP3271 | 2.5 mL |
| 1M citric acid, pH 6 | Fisher Scientific, BP327 and A142-212 | 90 µL |
| 10% Tween-20 | Sigma-Aldrich, P2287 | 100 µL |
| nuclease-free water | | to 10 mL final volume |
| **5X SSCT (store at -4°C for up to one month)** | | |
| Component | Source | Volume |
| 20X SSC | Fisher Scientific, S2713 and BP3271 | 2.5 mL |
| 10% Tween-20 | Sigma-Aldrich, P2287 | 100 µL |
| DEPC-treated water | Genesee Scientific, 20-138 | to 10 mL final volume |
| **Amplification buffer (pH 6.8, store at room temperature)** | | |
| Component | Source | Amount |
| sodium chloride | Fisher Scientific, S2713 | 292 mg |
| sodium phosphate dibasic | Mallinckrodt, 7917 | 71 mg |
| nuclease-free water | | to 10 mL final volume |

**Table S5.** Buffer recipes used in RNA-FISH. Volumes are given to make the buffers at 10 mL scale.

| Channel | Laser (nm) | Filter | PMT Voltage |
|---|---|---|---|
| tagBFP | 405 | 440 / 50 | 220 |
| Alexa Fluor™ 514 | 488 | 530 / 30 | 260 |
| mRuby2 | 561 | 620 / 15 | 260 |
| iRFP720 | 637 | 720 / 30 | 340 |
| FSC | | | 100 |
| SSC | | | 360 |

**Table S6.** Attune NxT flow cytometer channels and voltages

| | Alexa Fluor™ 514-A | mRuby2-A | tagBFP-A | iRFP720-A |
|---|---|---|---|---|
| Alexa Fluor™ 514-A | 1 | 0 | 0 | 0 |
| mRuby2-A | 0.01 | 1 | 0.015 | 0 |
| tagBFP-A | 0 | 0 | 1 | 0 |
| iRFP720-A | 0 | 0 | 0 | 1 |

**Table S7.** HCR Flow-FISH compensation matrix

| Gene | Forward primer | Reverse primer |
|---|---|---|
| mRuby2 | CCTTGAGGATGGCTGTCTCG | ATGGCCACCACCATCAACTT |
| GAPDH | GTATCGTGGAAGGACTCATGAC | ACCACCTTCTTGATGTCATCAT |

**Table S8.** Primer sequences used in RT-qPCR experiments
